# Structural basis for voltage-sensor trapping of the cardiac sodium channel by a deathstalker scorpion toxin

**DOI:** 10.1101/2020.12.28.424592

**Authors:** Daohua Jiang, Lige Tonggu, Tamer M. Gamal El-Din, Richard Banh, Régis Pomès, Ning Zheng, William A. Catterall

## Abstract

Voltage-gated sodium (NaV) channels initiate action potentials in excitable cells, and their function is altered by potent gating-modifier toxins. The α-toxin LqhIII from the deathstalker scorpion inhibits fast inactivation of cardiac NaV1.5 channels with IC50=11.4 nM. Here we reveal the structure of LqhIII bound to NaV1.5 at 3.3 Å resolution by cryo-EM. LqhIII anchors on top of voltage-sensing domain IV, wedged between the S1-S2 and S3-S4 linkers, which traps the gating charges of the S4 segment in a unique intermediate-activated state stabilized by four ion-pairs. This conformational change is propagated inward to weaken binding of the fast inactivation gate and favor opening the activation gate. However, these changes do not permit Na^+^ permeation, revealing why LqhIII slows inactivation of NaV channels but does not open them. Our results provide important insights into the structural basis for gating-modifier toxin binding, voltage-sensor trapping, and fast inactivation of NaV channels.

## Introduction

Eukaryotic voltage-gated sodium (NaV) channels generate the inward sodium current that is responsible for initiating and propagating action potentials in nerve and muscle ^1,2^. The sodium current is terminated within 1-2 milliseconds by fast inactivation ^1,2^. A wide variety of neurotoxins bind to six distinct receptor sites on NaV channels and modify their function ^3,4^. α-Scorpion toxins and sea anemone toxins bind to Neurotoxin Receptor Site 3, dramatically inhibit fast inactivation of NaV channels, and cause prolonged and/or repetitive action potentials ^3–5^. Scorpions utilize these toxins in their venoms to immobilize prey by inducing paralysis and causing cardiac arrhythmia ^4,6–8^. Because of their high affinity and specificity, scorpion toxins are used widely to study the structure and function of Na_V_ channels. α-Scorpion toxins bind to the voltage sensor (VS) in domain IV (*D*IV), which is important for triggering fast inactivation ^9–13^. Therefore, structures of the high-affinity complexes of α-scorpion toxins and NaV channels will provide critical information for understanding the structural basis for toxin binding, voltage-sensor trapping, and fast inactivation.

Eukaryotic NaV channels contain four homologous, nonidentical domains composed of six transmembrane segments (S1-S6), organized into a voltage-sensing module (VS, S1-S4) and a pore module (PM, S5-S6) with two intervening pore helices (P1 and P2) ^14,15^. The S4 segments contain four to eight repeats of a positively charged residue (usually Arg) flanked by two hydrophobic residues. These positively charged residues serve as gating charges, moving outward upon depolarization to initiate the process of activation ^14,15^. Chemical labeling and voltage clamp fluorometry suggest that *D*I-VS and *D*II-VS are primarily responsible for activation of the channel, whereas *D*IV-VS induces fast inactivation ^14,15^. A triple hydrophobic motif, Ile-Phe-Met (IFM), located in the *D*III-*D*IV linker, serves as the fast inactivation gate ^14,15^. Mutation of the IFM motif can completely abolish fast inactivation ^14,15^.

Determination of the structures of prokaryotic ^16–18^ and eukaryotic ^19–21^ Na_V_ channels has remarkably enriched our understanding of their structure and function. Those structures revealed that Na_V_ channels share similar key structural features^22^. The central pore is formed by the four PMs with the four VSs arranged in a pseudosymmetric square array on their periphery. The four homologous domains are organized in a domain-swapped manner, in which each VS interacts most closely with the PM of the neighboring domain. The four S6 segments come together at their intracellular ends to form the activation gate ^16–18^. Intriguingly, in the structures of mammalian NaVs, the IFM motif binds in a receptor site formed by the *D*III S4-S5 linker and the intracellular ends of the S5 and S6 segments in *D*IV, which suggests a local allosteric mechanism for fast inactivation of the pore by closing the intracellular activation gate ^19–21^.

The α-scorpion toxins bind to *D*IV-VS in its resting state, trap it in an intermediate activated conformation, and inhibit fast inactivation, providing an attractive target for studying the coupling of *D*IV-VS to pore opening and fast inactivation ^9–13,23,24^. Strong depolarization can reverse voltage-sensor trapping and drive the α-scorpion toxin off its receptor site, providing direct evidence for a toxin-induced conformation of the VS ^9,24,25^. The cryo-EM structure of the α-scorpion toxin AaHII was resolved bound to two different sites on a nonfunctional chimera of the cockroach sodium channel NaVPas, which contained 132 amino acid residues of the *D*IV-VS of the human neuronal sodium channel Na_V_1.7 embedded within 1449 residues of NaVPas ^26^. These results revealed structures of AaHII bound to the voltage sensors in both *D*I and *D*IV but did not resolve whether AaHII bound to either of these sites was functionally active in the chimera ^26^. Therefore, the precise structural mechanism by which α-scorpion toxin binds to the *D*IV-VS in a native sodium channel and blocks fast inactivation still remains elusive.

LqhIII from the ‘deathstalker scorpion’ *Leiurus quinquestriatus hebraeus* (also known as the Israeli yellow scorpion and the North African striped scorpion) is classified as an α-scorpion toxin and shares the common βαββ scaffold containing four pairs of Cys residues that form disulfide bonds ^7^. Most scorpion toxins paralyze prey by targeting the sodium channels in nerve and skeletal muscle specifically ^7^. In contrast, LqhIII binds with highest affinity to the human cardiac sodium channel, with an estimated *EC_50_* of 2.5 nM ^27,28^. It prevents fast inactivation efficiently, and it dissociates at an extremely slow rate ^27,28^, making it exceptionally potent.

In this work, we elucidate the molecular mechanisms of voltage-sensor trapping and block of fast inactivation by α-scorpion toxins in the context of a functional native toxin-receptor complex by determining the cryo-EM structure of rat cardiac sodium channel Na_V_1.5 in complex with the α-scorpion toxin LqhIII at 3.3 Å resolution. Our experiments provide important insights into the structural basis for gating-modifier toxin interaction, voltage-sensor trapping, electromechanical coupling in the VS, and fast inactivation of the pore.

## Results

### Voltage-sensor trapping of Na_V_1.5 by LqhIII

For our structural studies, we took advantage of the fully functional core construct of the rat cardiac sodium channel NaV1.5 (rNa_V_1.5_C_), which can be isolated with high yield and high stability ^21^. Expression of rNaV1.5C in the human embryonic kidney cell line HEK293S GnTI^−^ and recording from single cells in whole-cell patch clamp mode (see Methods) yields inward sodium currents that activate rapidly and inactivate within 6 ms (Fig. 1a, black trace; inward current is plotted as a negative quantity by convention). Perfusion of increasing concentrations of LqhIII progressively slows the fast inactivation process and makes it incomplete (Fig. 1a, colored traces). We measured the sodium current remaining 6 ms after the depolarizing pulse as a metric of LqhIII toxin action (Fig. 1a, dotted line), because the unmodified sodium current has declined to nearly zero by this time, whereas substantial toxin-modified sodium current remains. The EC_50_ value for the increase in sodium current remaining at 6 ms following the stimulus is 11.4 nM (Fig. 1a). This effect of LqhIII and other α-scorpion toxins is achieved by trapping the voltage sensor in *D*IV of sodium channels in a conformation that allows sodium channel activation but prevents coupling to fast inactivation ^4,9,10^. Voltage-sensor trapping develops slowly and progressively over more than 20 min, with a half-time of 11.3 min at 100 nM (Fig. 1b). As expected from previous work ^4,9,10^, strong depolarizing pulses to +100 mV cause dissociation of the toxin and loss of its blocking effect on fast inactivation (Fig. 1c). The molecular mechanism for this long-lasting voltage-dependent block of fast inactivation of NaV1.5 sodium currents by LqhIII is unknown.

**Fig. 1.**
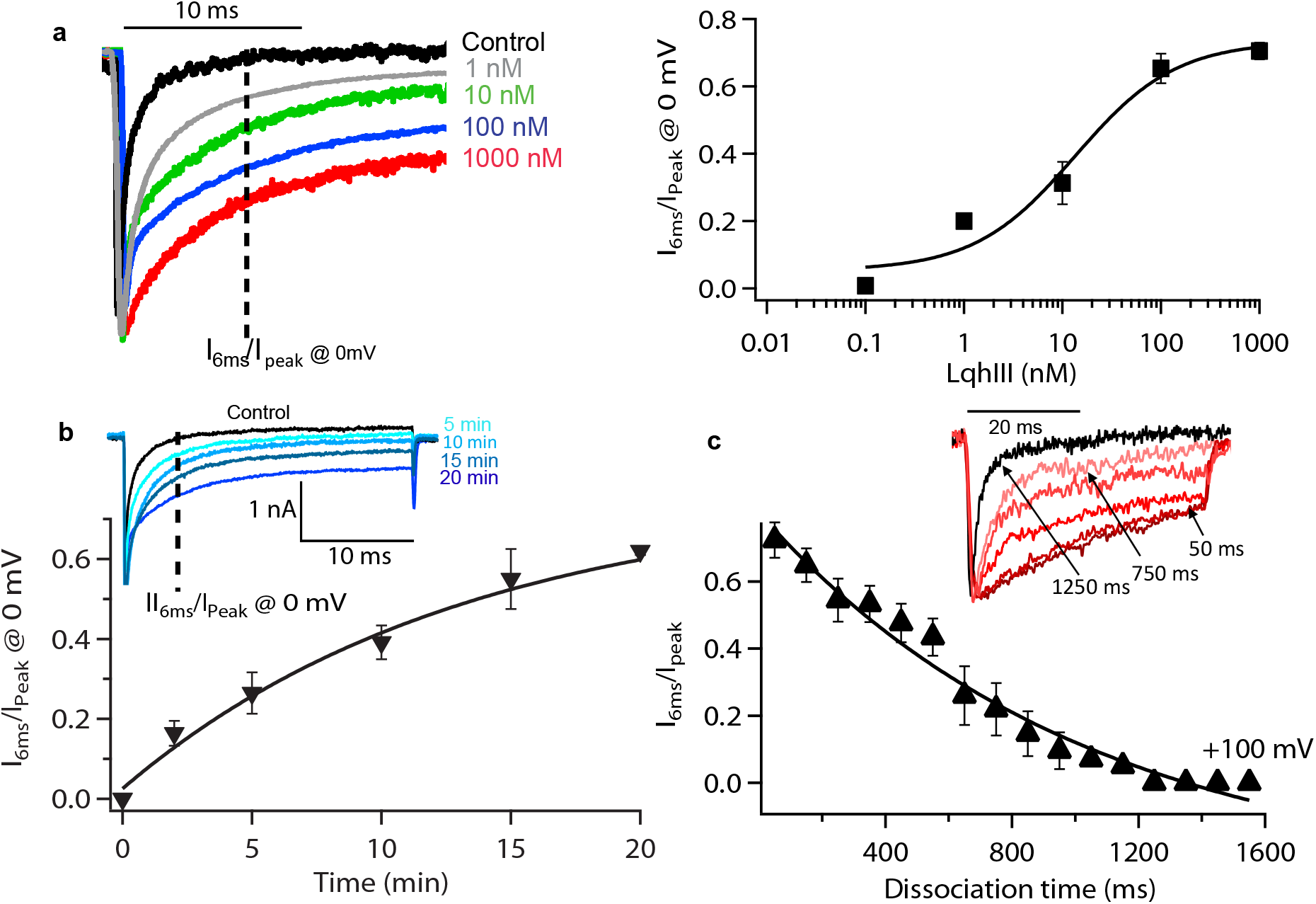
Block of fast inactivation of rNa_V_1.5_C_ by LqhIII. **a**. *Left.* Normalized current traces from HEK293 cells expressing rNa_V_1.5_C_ in the absence (black) or in the presence of 1 nM (grey), 10 nM (green), 100 nM (blue), or 1000 nM (red) LqhIII. Cells were held at −120 mV and Na^+^ currents were elicited with a 1000-ms step to 0 mV. Measurements at different toxin concentrations were carried out on different cells because of the limited stability of the whole-cell recording configuration on virus-infected HEK cells. *Right.* Concentration-response curve of LqhIII scorpion toxin for inhibition of fast inactivation of the rNa_V_1.5_C_ channel. Each point is an average of 4-5 cells. Data points error bars represent mean and s.e.m. The solid line represents the Hill equation fit to the data. EC_50_ = 11.4 ± 0.9 nM (s. e. m., n = 23 cells). **b**. The time course of association of 100 nM of LqhIII scorpion toxin. Cells were held at −120 mV and the toxin was perfused. A pulse to 0 mV from V_m_ = −120 mV was applied at the indicated times. Single exponential fitting of the block of inhibition ratio showed a time constant of 11.3 min. Each point is an average of 6 cells. Data points and error bars represent mean and s.e.m. **c.**Time course of LqhIII dissociation. A three-pulse protocol was applied: first, a pulse from −120 mV to +100 mV for the indicated times, followed by a second 50-ms hyperpolarizing pulse to allow recovery from fast inactivation, and finally by a third pulse of 50 ms to 0 mV to measure the extent of recovery of fast inactivation kinetics. Mean and s. e. m.; n = 5 cells for each data point. *Inset,* representative traces showing recovery of fast inactivation.

### Structure determination of rNa_V_1.5_C_/LqhIII complex by cryo-EM

We analyzed the structural basis for the potent voltage-sensor trapping effects of LqhIII by cryogenic electron microscopy (cryo-EM). LqhIII was incubated with purified rNa_V_1.5_C_ for 30 min. The regulatory proteins FGF12b and calmodulin were added to stabilize the isolated protein, and the toxin/channel complex was further purified by size-exclusion chromatography (SEC). A symmetric peak of the toxin/channel complex was collected from the second SEC run (Supplementary Fig. 1a, b). Detailed descriptions of protein expression, purification, cryo-EM imaging, and data processing are presented in Methods.

Cryo-EM data were collected on a Titan-Krios electron microscope and processed using RELION (Supplementary Fig. 1c, d; Supplementary Fig. 2a-c). A 3D reconstruction map of the rNa_V_1.5_C_/LqhIII complex was obtained at an overall resolution of 3.3 Å, based on the Fourier Shell Correlation (FSC) between independently refined half-maps (Fig. 2a). Strong density specifically localized near the extracellular side of *D*IV-VS shows that there is only one LqhIII molecule bound to rNa_V_1.5_C_ (Fig. 2b; purple), as expected from previous biochemical studies of scorpion toxin binding to sodium channels ^9^. The local resolution for the PM core region is ~3.0-3.5 Å, whereas the four peripheral VSs have local resolutions of ~3.5-4.0 Å (Fig. 2c). The resolution for the toxin is lower than the channel protein (~4.0-5.0 Å, Fig. 2c). However, the interacting surface of the toxin that binds to *D*IV-VS has a resolution of ~3.5-4.0 Å for the amino acid side chains that form the complex, as they are tightly bound (Supplementary Fig. 2d and e). The 3D structure of the tightly disulfide-crosslinked toxin is well-known from previous studies (Supplementary Fig. 3a) ^29^, allowing it to be accurately fit into the observed density. No significant density was observed at high resolution for the C-terminal domain (CTD), FGF12b, or calmodulin (Fig. 2b), indicating that these components of the purified protein complex are mobile.

**Fig. 2.**
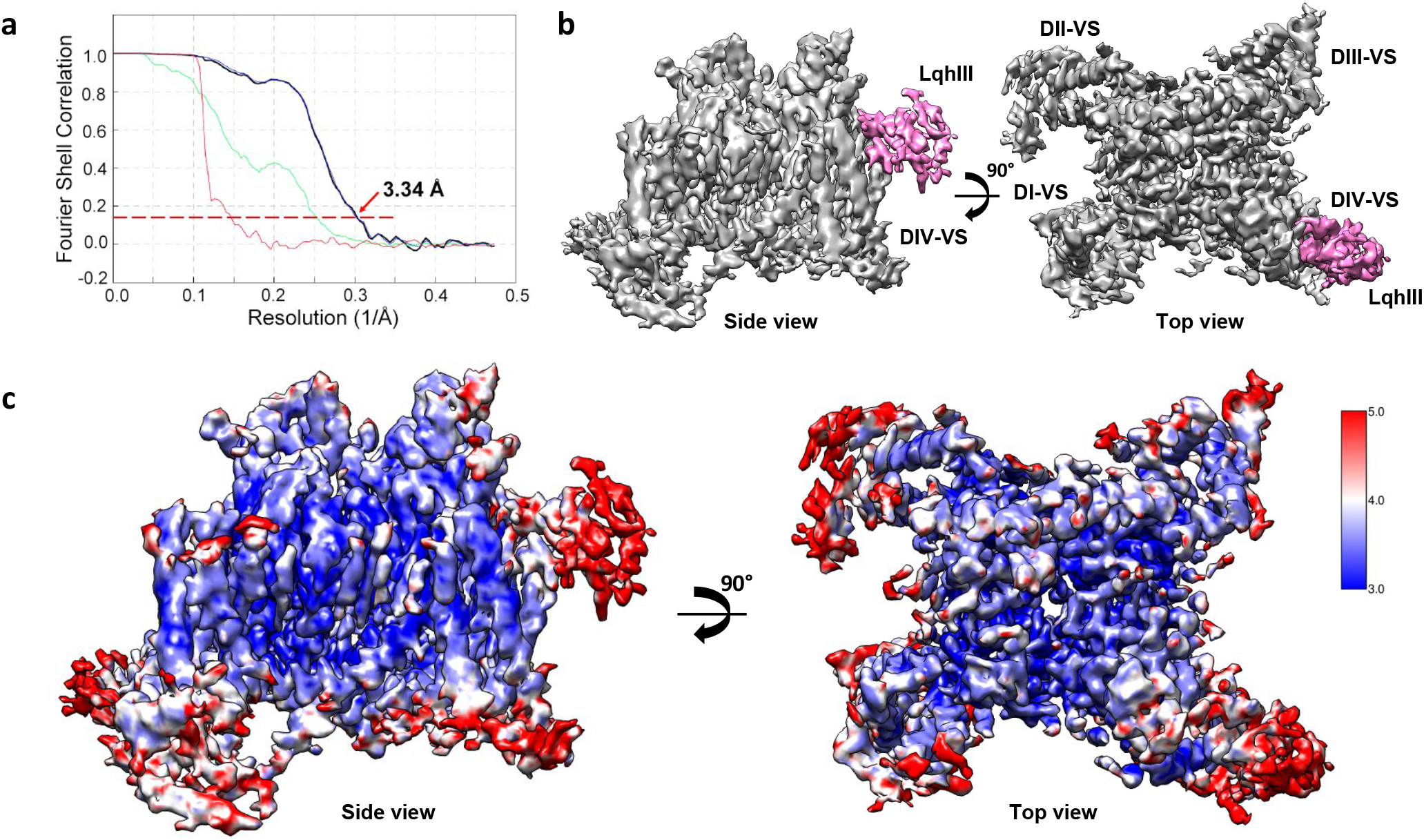
Cryo-EM structure of the rNa_V_1.5_C_/LqhIII complex. **a.**The FSC between independently refined half-maps for rNa_V_1.5_C_/LqhIII reconstruction. **b**. Overall cryo-EM reconstruction (side view, left; top view, right) of the rNa_V_1.5_C_/LqhIII complex. Na_V_1.5_C_ and LqhIII colored in grey and purple, respectively. **c**. Local resolution (side view, left; top view, right) of the EM map colored from blue to red representing resolution from high to low (side bar).

### Overall structure and LqhIII binding site

The high-resolution cryo-EM density map allowed us to build an atomic model for the rNa_V_1.5_C_/LqhIII complex (Fig. 3a and b). The overall structure of the rNa_V_1.5_C_/LqhIII complex is very similar to our previous apo-rNa_V_1.5_C_ structure ^21^, with a minimum RMSD of 0.78 Å over 1164 residues. However, local conformational differences give many important insights. The structure of LqhIII is rigidly locked by disulfide bonds, except for the β2β3 loop and C-terminal region, which are highly flexible in solution as revealed by NMR analyses (Fig. 3c). Remarkably, LqhIII uses these two flexible regions to bind to the extracellular side of *D*IV-VS by wedging its β2β3 loop and C-terminus into the aqueous cleft formed by the S1-S2 and S3-S4 helical hairpins (Fig. 3b). These features are in close agreement with previous molecular-mapping studies of neurotoxin receptor site 3 ^11^ and with the structure of the AaHII/Na_V_Pas-Na_V_1.7 chimera ^26^ (see Discussion). The toxin may attack Neurotoxin Receptor Site 3 in the *D*IV-VS using its most flexible regions to allow it to dock in a stepwise manner that results in a tight induced-fit complex.

**Fig. 3.**
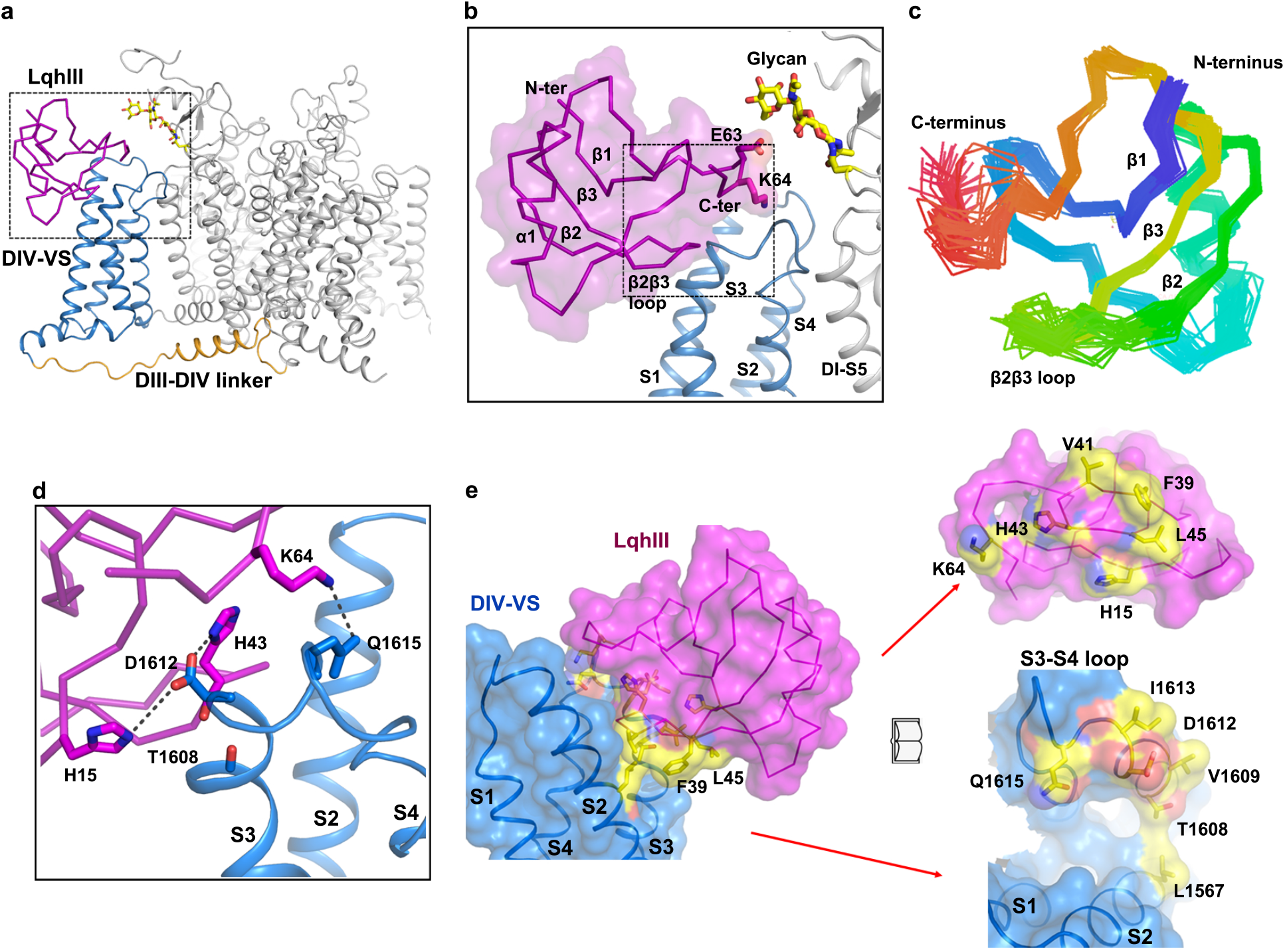
Overall structure of rNa_V_1.5_C_/LqhIII complex and LqhIII binding site. **a.**Cartoon representation of the overall structure of the Na_V_1.5_C_/LqhIII. LqhIII, *D*IV-VS and *D*III-*D*IV linker were colored in purple, blue and orange, respectively. The same color scheme is applied hereafter in this manuscript unless specified otherwise. The glycosyl moieties shown in sticks colored in yellow. The black dash square is indicated for panel b. **b**. Zoom-in view of LqhIII binding to the *D*IV-VS. The black dash square indicated for panel d. **c.** The NMR structure of the LqhIII (PDB code: 1FH3) indicating the flexibility of the β2β3 loop and C-terminus. **d.** Detailed interactions between LqhIII and *D*IV-VS. Key residues shown in sticks were labeled. Interaction surfaces of the *D*IV-VS (blue) and the LqhIII (purple). Key residues for the interaction shown in yellow shading and embedded sticks.

The close interactions of the C-terminus and the β2β3 loop of LqhIII with rNa_V_1.5_C_ are illustrated in Fig. 3b and d. At the C-terminus of LqhIII, Glu63 interacts with the Asn329-linked glycan from *D*I-PM, and Lys64 dips into the aqueous cleft and interacts with Gln1615 (Fig. 3b and d). The end of the β2β3 loop inserts into the *D*IV-VS cleft and partially unwinds the last helical turn of the S3 segment. Mutagenesis studies mapping Neurotoxin Receptor Site 3 revealed a negatively charged residue in the extracellular S3-S4 linker that is conserved among Na_V_ channels and is critical for α-scorpion toxin binding ^4,9^. In agreement with those studies, the conserved negatively charged residue Asp1612 at an equivalent position in Na_V_1.5 mediates this crucial interaction with the bound toxin. His43 and His15 wrap around Asp1612 like pincers forming a hydrogen bond (~2.5 Å) and a potential salt bridge (~4.0 Å), respectively (Fig. 3d). Moreover, we note that the backbone carbonyl of His43 engages the backbone carbonyl of Thr1608 at a distance of 2.8-3.5 Å, which may contribute to the affinity or specificity of interactions with the β2β3 loop (Fig. 3d)^30^.

The complementary interacting surfaces of LqhIII and Neurotoxin Receptor Site 3 are depicted in a space-filling model in Fig. 3e (left), and the functionally important interacting residues are highlighted in yellow with embedded sticks and displayed in an ‘open-book’ format in Fig. 3e (right). The interacting surface area of neurotoxin receptor site 3 covers ~ 836 Å^2^ located on an arc stretching from the S3-S4 linker to the S1-S2 linker (Fig. 3e, right). The LqhIII toxin latches onto that arc, gripping it between the β2β3 loop and the C-terminus (Fig. 3e). It is likely that the flexibility of these regions of the toxin in solution is important for its initial approach and final tight grip on its target site.

### An intermediate activated state of*D*IV-VS trapped by LqhIII

Fast inactivation of Na_V_ channels requires activation of *D*IV-VS ^9,10,12,13^. Because there is no membrane potential during solubilization and purification, the VSs of published Na_V_ structures are usually in partially or fully activated states. In our apo-rNa_V_1.5_C_ structure, four of the six gating charges of *D*IV-VS pointed outward on the extracellular side of the hydrophobic constriction site (HCS), as expected for an activated state ^21^. As a result, the fast inactivation gate in the apo-rNa_V_1.5_C_ structure binds tightly in a hydrophobic pocket next to the activation gate ^21^. α-Scorpion toxins bind to Na_V_ channels in the resting state with higher affinity and trap the channel in a partially activated state, in which both the rate and extent of transition to the inactivated state are impaired (Fig.1) ^9,10^. Because of its high affinity and specificity, LqhIII is able bind to the purified rNa_V_1.5_C_ protein in its activated state and chemically induce voltage-dependent structural changes to partially deactivate the VS. Remarkably, LqhIII binding drives *D*IV-S4 approximately two helical turns inward to form an intermediate, partially activated structure (Fig. 4a and b). Each gating charge Arg in the intermediate activated *D*IV-VS is positioned ~10-12 Å further inward than in the fully activated *D*IV-VS (Fig. 4a and b). Importantly, in the toxin-bound intermediate activated state reported here, R1 to R4 adopt a 3_10_-helix conformation, with the last helical turn of the S4 segment relaxing R5 into alpha-helical form. In contrast, in the fully activated state, the region between R2 to R6 is in 310-helical form, but R1 is alpha-helical. As a consequence of the 3_10_-helix conformation from R1-R4 in the toxin/channel complex, the residues between R1-R2 and R3-R4 bridge the HCS such that R1-R2 and R3-R4 share the same vertical plane in their interactions with the negative side chains of the extracellular negative cluster (ENC) and intracellular negative cluster (INC), respectively. This unique linear voltage-sensor-trapped conformation would be strongly stabilized by these simultaneous gating charge interactions outside and inside the HCS, which may provide the chemical energy required for potent voltage-sensor trapping against the force of the transmembrane electrical field and therefore for effective modification of sodium channel gating. The potential gating charges R5 and R6 translocate to the intracellular side of the VS completely. These charged residues were proposed to interact with the CTD in the structure of the Na_V_Pas/Na_V_1.7 chimera ^26^. However, the CTD was not resolved in our structure, preventing visualization of the potential binding positions of R5 and R6.

**Fig. 4.**
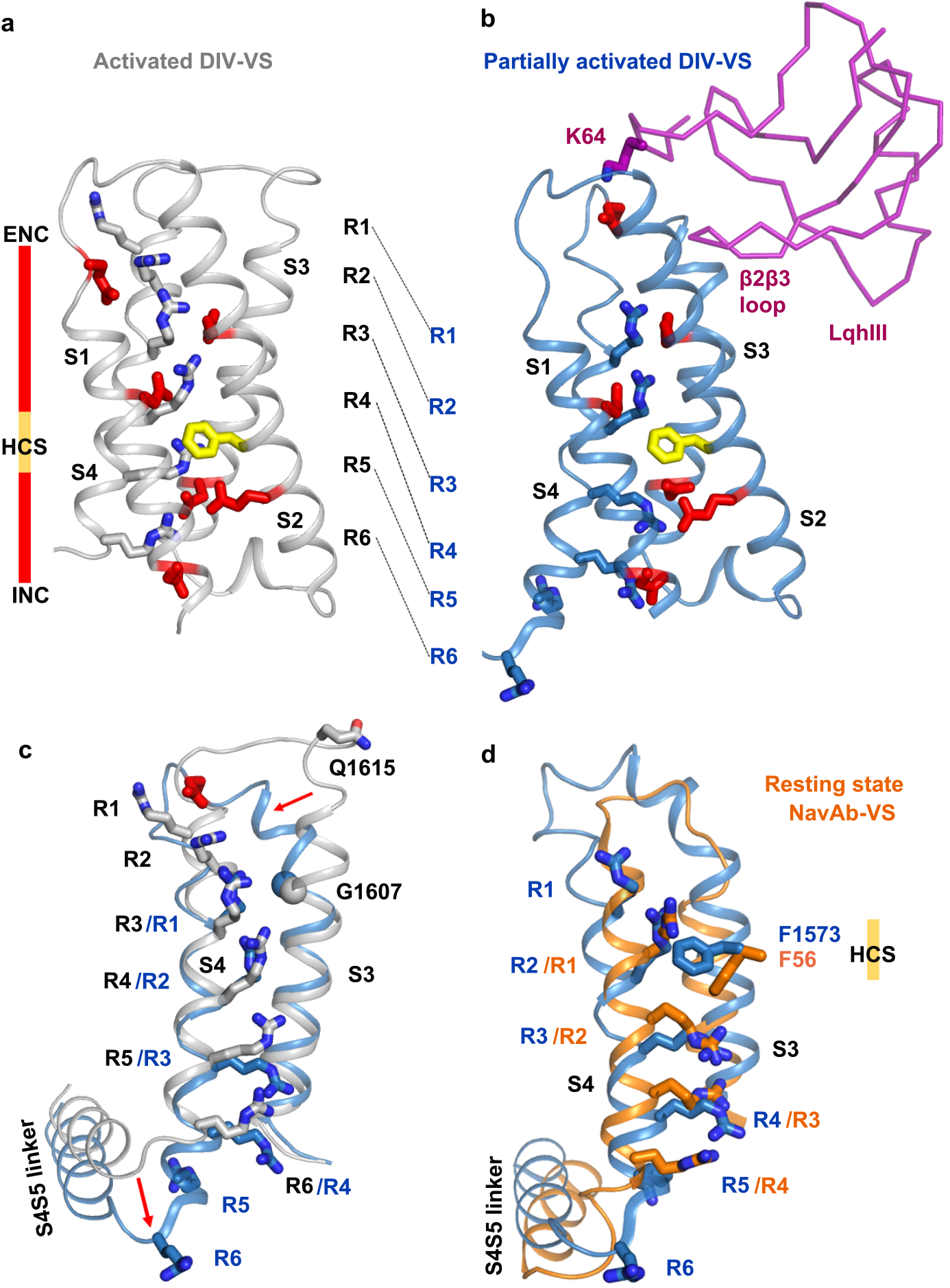
Conformational Change of *D*IV-VS. **a, b**. Structures of the activated Na_V_1.5 *D*IV-VS and intermediate activated Na_V_1.5 *D*IV-VS were colored in grey and blue respectively. Gating charges (grey or blue), ENC (red), HCS (yellow) and INC (red) were shown in sticks. Shift of each gating charge was indicated by black dashed lines. **c.** Superposition of Na_V_1.5 *D*IV-VS between activated and intermediate activated state. Red arrows indicate the conformational changes. **d** Superposition of the intermediate activated Na_V_1.5 *D*IV-VS and resting state Na_V_Ab-VS.

Superposition of the fully activated state (grey) and toxin-induced intermediate activated state (blue) of the *D*IV-VS revealed a remarkable conformational difference (Fig. 4c). From S1 through most of S3 there is little or no structural change, whereas the final two helical turns of S3 and the entire S4 segment undergo dramatic conformational shifts. Notably, Gly1607 serves as a pivot point for S3 rotation, and the rotation of upper S3 in turn moves S4 downward ~11 Å, such that R1 and R2 in the intermediate activated state are approximately in the positions of R3 and R4 in the fully activated state (Fig. 4c). This toxin-induced conformational change in the S3-S4 linker is further documented by our fit to the cryo-EM density, which is illustrated in Extended Fig. 3c-e. At the intracellular end of S4, an elbow-like bend is formed between S4 and the S4-S5 linker, which pushes the S4-S5 linker ~4.6 Å inward at its N-terminal end (Fig. 4c). Intriguingly, our previous resting-state structure of Na_V_Ab showed that a similar elbow pushes the S4-S5 linker and its connection to S4 strikingly inward and twists this segment in order to close the intracellular activation gate ^31^. This conformational change in the S4-S5 linker is further supported by the close fit of our structural model to the cryo-EM density (Supplementary Fig. 4a). Superposition of the intermediate activated *D*IV-VS structure (blue) upon the resting state Na_V_Ab-VS structure (orange) further illuminates these conformational differences (Fig. 4d). The connecting S3-S4 loop of the intermediate activated state of the LqhIII/rNa_V_1.5_C_ complex is not located as deeply inward as that of resting state of Na_V_Ab and is not as tightly twisted (Fig. 4d). Moreover, the R1 and R2 gating charges are both located fully outward from the HCS in the partially activated S4 segment in the LqhIII/rNa_V_1.5_C_ complex, whereas R1 is positioned only partially outward from the HCS in the resting state of Na_V_Ab (Fig. 4d). In addition, the S4-S5 linker in the intermediate activated state has not moved as deeply into the cytosol as in the resting state (Fig. 4d). These differences suggest that the toxin-induced intermediate activated state of Na_V_1.5 VS is indeed an intermediate state between the resting state and the fully activated state.

A hallmark feature of the action of α-scorpion toxins is strongly voltage-dependent dissociation from their receptor site, which correlates with the voltage dependence of activation of sodium channels (Fig. 1c; ^9,24,25^). The structure of the rNa_V_1.5_C_ /LqhIII toxin complex reveals the molecular basis for this important aspect of scorpion toxin action. In the complex of the toxin with the partially activated state of the *D*IV VS, the positive charge of the ɛ-amino group of K64 on LqhIII interacts with the same negatively charged side chain in the ENC that interacts with R1 and R2 in the activated conformation of the VS (Fig. 4a and b). In light of these structures, it seems likely that outward movement of the S4 segment during activation of the *D*IV VS creates a clash with K64 and causes toxin dissociation by both electrostatic repulsion and steric hindrance. This potent combination of electrostatic repulsion and physical clash is sufficient to overcome the high binding energy of the α-scorpion toxins.

### Partially open intracellular activation gate

Rapid voltage-dependent activation is one of the signature functions of Na_V_ channels ^1,2^. Full understanding of the mechanism of activation requires structural information on Na_V_ channels in different states. Side-by-side comparison of our previous activated-state structure with our current LqhIII-bound intermediate activated structure of rNa_V_1.5_C_ reveals key steps in the coupling of the conformational changes of the *D*IV-VS to the intracellular activation gate (Fig. 4c and 5). To reach the intermediate activated state (blue) from the fully activated state (grey), S4 of *D*IV-VS moves inward ~11 Å and pushes the *D*IV S4-S5 linker inward ~4.6 Å through formation of an elbow-like bend (Fig. 4c and 5a). The shifted *D*IV S4-S5 linker engages *D*IV-S6 through Ser1655 (Fig. 4c and 5c). As a result, *D*IV-S6 shifts toward the pore axis, and this movement pushes *D*I-S6 away from the center of the orifice (Fig. 5a, b and Supplementary Fig. 4b and c). Meanwhile, the *D*I S4-S5 linker moves outward away from the activation gate, which shifts both *D*I-S6 and *D*II-S6 away from pore axis and contributes to a more open conformation. The shifts of the four S6 segments result in an enlarged activation gate with a van der Waals diameter of 6.6 Å (Fig. 5b and Supplementary Fig. 4b and c), which is ~1 Å larger than the activation gate in apo-rNa_V_1.5_C_, but ~2 Å smaller than the expected orifice of the activation gate of fully open rNa_V_1.5_C_ when modeled using the open state of Na_V_Ab ^21,32^. Unambiguous density shows Tyr1769 in *D*IV-S6 in an outward-pointing conformation making the opening of the activation gate larger, whereas it is pointed inward in the fully activated structure (Fig. 5b). Together, these conformational movements result in a wider opening in the activation gate in the toxin-induced intermediate activated state, which could in principle be sufficient for sodium conductance.

**Fig. 5.**
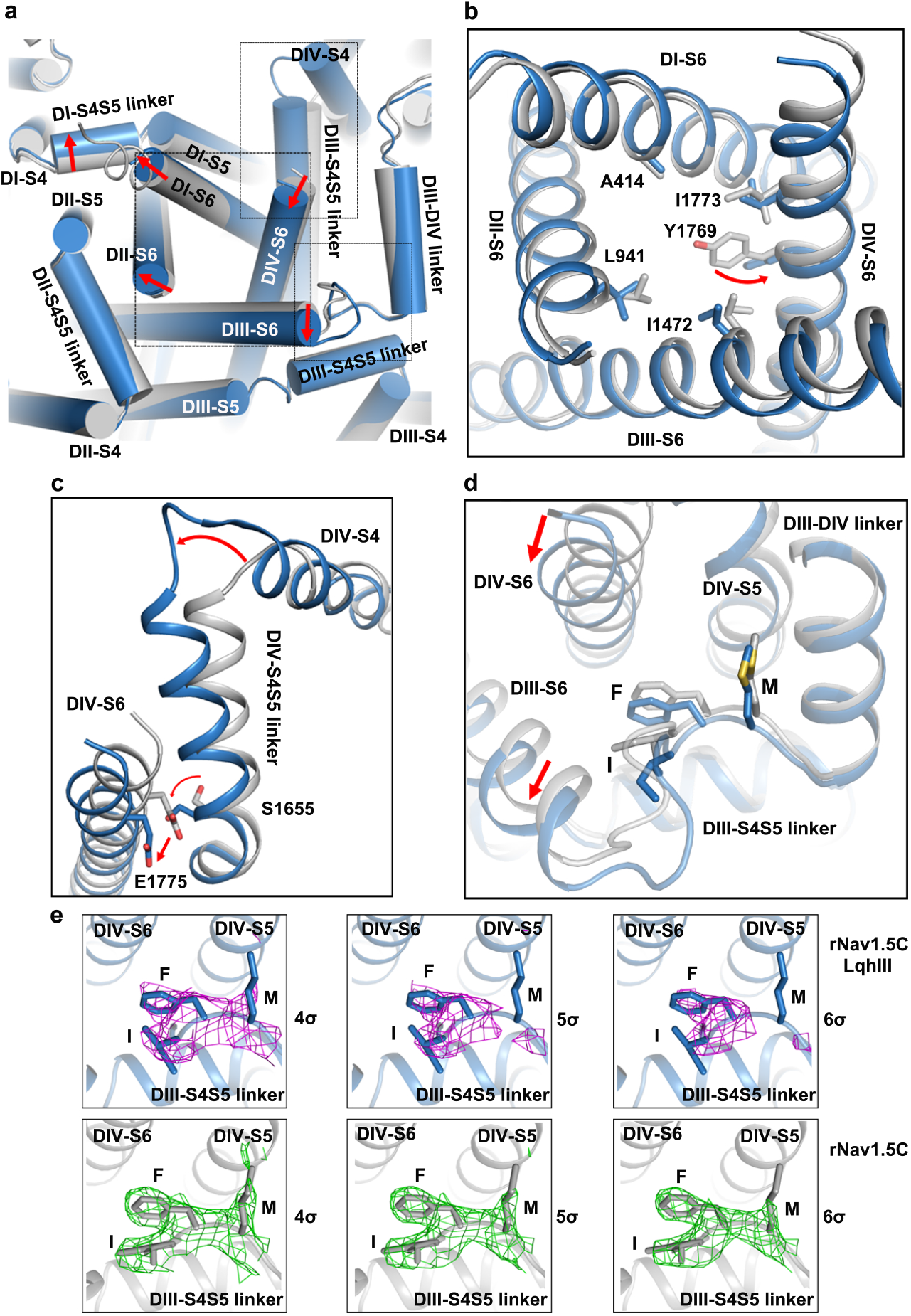
Coupling of LqhIII binding and conformational change in *D*IV-VS to the S4-S5 linkers and the intracellular activation gate. **a.** Superposition of Na_V_1.5 intracellular gate between activated (grey) and intermediate activated (blue) states of the *D*IV-VS. Red arrows indicate the conformational changes. The black dash squares indicated for panel b-d. **b.** Zoom-in view of the intracellular activation gate with constriction residues shown in sticks. **c**. Zoom-in view of *D*IV S4-S5 linker mediates the conformational changes. **d.** Zoom-in view of IFM motif shift between activated and deactivated state. e. Close-up views of the IFM motif in the free and LqhIII-complexed Na_V_1.5 together with its cryo-EM density contoured at different σ levels.

Consistent with the more open conformation of the activation gate, we observed much stronger density of bound lipid or detergent in the lumen of the activation gate in our intermediate activated rNa_V_1.5_C_ structure compared to that of the activated apo-rNa_V_1.5_C_ structure (Supplementary Fig. 4b and c). Considering that the two proteins were purified following the same procedures, it is likely that the same set of lipid and detergent molecules would be available for binding. Therefore, we believe the larger lipid or detergent molecule is able to bind in the lumen of the activation gate of the intermediate activated structure because of its larger diameter rather than because of a change in lipid or detergent concentrations between the two protein preparations.

### Loosely bound fast inactivation gate

In activated Na_V_ structures, the IFM motif of the fast inactivation gate binds to a hydrophobic pocket formed by *D*III S4-S5 linker and the intracellular ends of *D*IV-S5 and *D*IV-S6 ^19-21^. In the intermediate activated rNa_V_1.5_C_/LqhIII structure, weak, yet consistent, density for the IFM motif suggests that the conformational changes of the VS and activation gate have in turn altered the conformation of the bound IFM motif and made it less stable (Fig. 5d, e, Supplementary Fig. 4b and c). Evidently, the movement of *D*IV-S6 squeezes the binding pocket, and the shift of *D*III-S6 destabilizes binding of the IFM motif. We observed a small shift in the position of the IFM motif in the intermediate activated structure, as well as weaker density for the Met side chain, consistent with greater mobility and partial dissociation (Fig. 5d, e). These changes are illustrated most clearly in Fig. 5e by comparing cryo-EM density contoured at 4σ, 5σ, and 6σ in the presence and absence of LqhIII. We propose that α-scorpion toxins inhibit fast inactivation by both trapping the *D*IV-VS in an intermediate activated conformation and altering the shape of the IFM binding pocket, which causes slowed IFM binding, more rapid dissociation of the IFM motif, and release of the fast inactivation gate. These effects are responsible for the slowed rate of fast inactivation and the incomplete extent of fast inactivation that are hallmarks of the action of LqhIII and other α-scorpion toxins (Fig. 1).

### LqhIII does not open the intracellular activation gate

Our static cryo-EM view of the intracellular activation gate of apo-rNa_V_1.5_C_ suggests that it is partially open compared to our tightly closed resting-state model of rNa_V_1.5_C_ based on the Na_V_Ab resting state ^21,31^. However, it does not appear to be open enough to conduct hydrated Na^+^ ^32,33^. To test this hypothesis, we used molecular dynamics methods similar to those we previously applied to the Na_V_Ab structure ^32,33^ in order to investigate the effect of LqhIII on pore hydration and dilation of the intracellular activation gate (Fig. 6). The inner pore of rNa_V_1.5_C_ is depicted lying from right (extracellular) to left (intracellular) with the surrounding S5 and S6 helices illustrated in orange (Fig. 6a). Water molecules (red) fill the inner part of the central cavity on the right and the intracellular exit from the pore on the left. However, in this snapshot, there is a gap in hydration in the intracellular activation gate itself (white), where the S6 segments come together in a bundle (orange helices, Fig. 6a). In fact, statistical analysis of the conformational ensemble shows that the average probability density of water molecules in the intracellular activation gate (purple band) is near zero in simulations of both rNa_V_1.5_C_ (black) and rNa_V_1.5_C_/LqhIII (red; Fig. 6b). Accordingly, Na^+^ did not permeate through the dehydrated activation gate in any of the simulations, suggesting that the pore is functionally closed. Not only is the activation gate the least hydrated region of the pore on average, but it is also nearly always dehydrated (Fig. 6a and b). Even when a pathway connecting the central cavity to the intracellular space is transiently present, water molecules are usually excluded from entering this region due to the hydrophobic effect (Fig. 6a). As such, the activation gate is predominantly dehydrated (dewetted) and occupied by 4 water molecules on average, compared to 10-11 molecules of water in the open activation gate of Na_V_Ab (Lenaeus et al., 2017). Thus, the intracellular activation gate of r, much like that of other voltage-gated ion channels, fits the paradigm of a hydrophobic activation gate, where small increases in the size of the gate tilt the wetted/dewetted equilibrium towards the wetted state as a prerequisite to ion permeation ^32,34–37^. These results showing that the intracellular activation gate is functionally closed in the rNa_V_1.5_C_/LqhIII complex illustrate the structural basis for the well-established effect of α-scorpion toxins to slow and prevent fast inactivation of sodium channels without opening the pore and allowing sodium conductance.

**Fig. 6.**
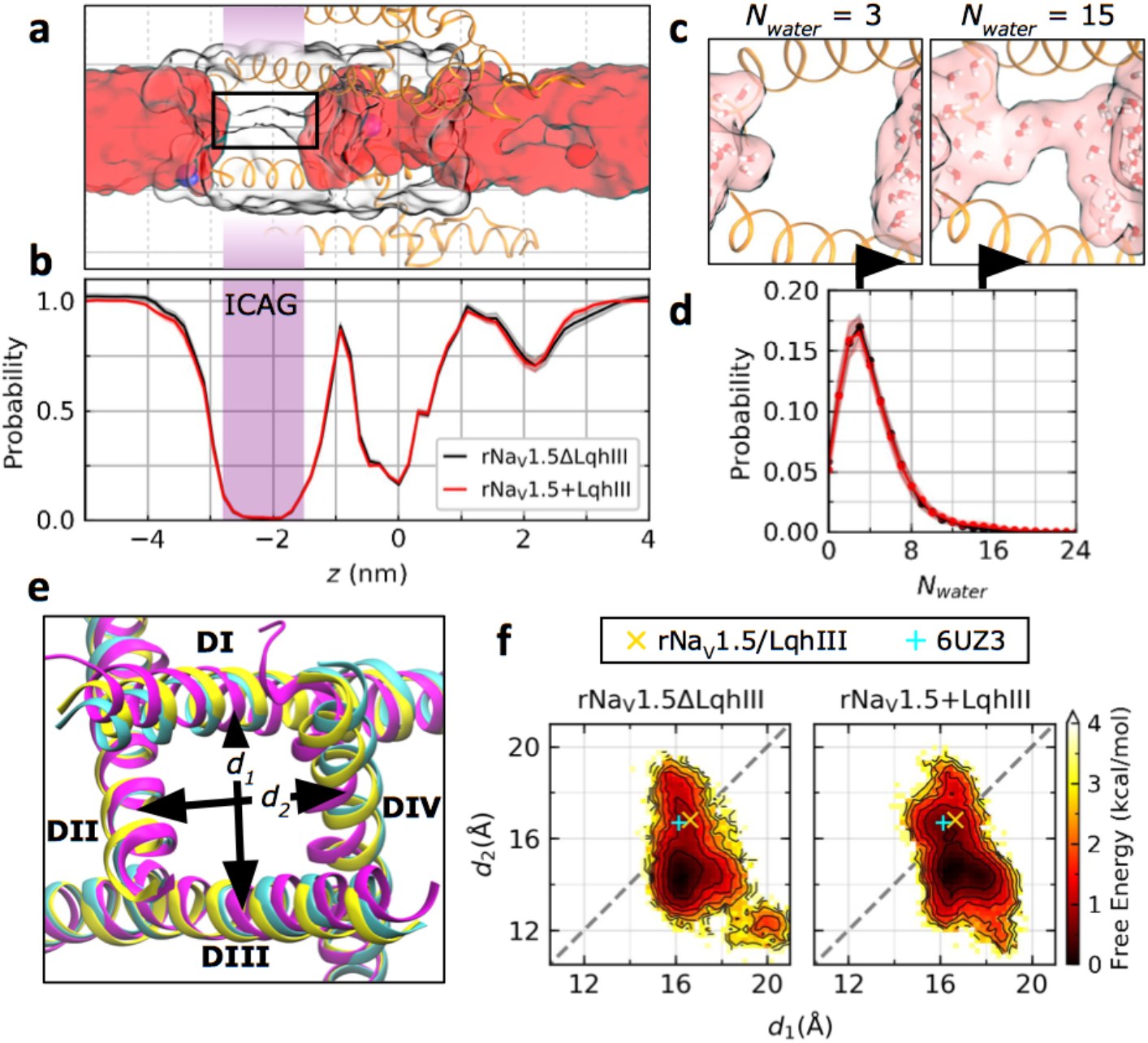
Molecular dynamics analysis of hydration and Na^+^ permeation through the rNa_V_1.5_C_/LqhIII complex. **a.** Side view of rNa_V_1.5_C_ (orange ribbons; domains II and IV) from MD simulations highlighting Na^+^ ions (blue spheres), the water-occupied volume within a cylinder of radius 8.5 Å (red surface), and the protein-occupied volume within a cylinder of radius 12 Å (colorless surface). In this snapshot, the protein cavity (outlined in black rectangle) at the intracellular activation gate (ICAG; purple shaded region, −2.8 nm < z < −1.5 nm) is dehydrated. The QuickSurf representation in VMD was used for surfaces. **b.** Average hydration along the pore-axis for simulations of rNa_V_1.5_C_/LqhIII (red line) with and (black line) without the toxin. Overall pore hydration was unchanged whether or not the toxin is included in simulations. Shading corresponds to standard error of the mean (s.e.m.). **c.** Molecular representations of the gate containing *N_water_*= 3 (left) or 15 (right) water molecules. **d**. Average probability distribution of *N_water_* in the gate. The gate is more likely to contain 10 or more water molecules when the toxin is present. Data points and shading represent mean and s.e.m. **e.** Bottom (intracellular) view of the activation gate for cryo-EM rNa_V_1.5_C_ (PDB ID: 6UZ3; cyan), cryo-EM rNa_V_1.5_C_/LqhIII (yellow), and an example conformation from the most sampled basin in MD (magenta) are superimposed. The distances *d_1_*, *d_2_* between opposing helices are shown schematically (see Methods). **f.** Free energy of *d_1_* vs. *d_2_* computed from MD simulations. The reference cryo-EM structures of (cyan +) rNa_V_1.5_C_ and (yellow ×) rNa_V_1.5_C_/LqhIII are indicated. Contour lines are shown every 0.5 kcal/mol from 0 to 4 kcal/mol. In the simulations, the intracellular activation gate often contracts and adopts an asymmetric conformational basin with *d_2_* < *d_1_*, with the symmetric conformation observed by cryo-EM corresponding to a metastable state 0.5 to 1 kcal/mol higher in free energy. Time frames spread across 30 independent simulations of rNa_V_1.5_C_ with or without LqhIII were used for analyses (n=29,026 frames for rNa_V_1.5_C_ and n=29,899 frames for rNa_V_1.5_C_/LqhIII).

### Fluctuations of the diameter of the activation gate

The distances between opposing S6 helix tails (*d_1_*: *D*I-*D*III, *d_2_*: *D*II-*D*IV) were monitored to estimate the dilation of the gate (Fig. 6e and f). The activation gate in the cryo-EM structure of rNa_V_1.5_C_/LqhIII is slightly wider and more symmetric [(*d1*, *d2*) = (16.6 Å, 16.8 Å)] than that of the cryo-EM structure of rNa_V_1.5_C_ [(*d_1_, d_2_*) = (16.1 Å,16.6 Å); Fig. 6e and f] ^21^. In the simulations, the structure of the gate fluctuates, with *d_2_* deviating by up to 4 Å from the toxin-bound cryo-EM structure (Fig. 6f). Overall, asymmetric conformations in which *d_2_* < *d_1_* are slightly favored (by 0.5 to 1.0 kcal/mol) relative to the symmetric cryo-EM conformation of rNa_V_1.5_C_/LqhIII. The average values of both *d_1_* and *d_2_* undergo small but significant increases relative to simulations without LqhIII, as does the average number of water molecules in the gate (Supplementary Figs. 5a). Although the pore remains predominantly dewetted irrespective of the presence of LqhIII, fluctuations leading to more dilated conformations of the activation gate are correlated with larger hydration numbers (Supplementary Fig. 6a), with transient occurrences of 10 or more water molecules connecting the central cavity to the intracellular environment (Fig. 6c and d). However, the activation gate is not significantly more likely to be wetted in simulations of rNa_V_1.5_C_ with LqhIII than in simulations without LqhIII (Supplementary Fig. 5b), consistent with the fact that toxin binding does not open the gate sufficiently for passage of Na^+^. Nevertheless, the analysis of fluctuations in diameter and hydration of the intracellular activation gate provide an initial suggestion that LqhIII binding may facilitate the transition to the open state of Na_V_1.5, which requires activation of the voltage sensors in domains I, II, and III for completion.

## Discussion

We determined the structure of rat Na_V_1.5C in complex with the α-scorpion toxin LqhIII by single particle cryo-EM. Biochemical and biophysical studies support only a single neurotoxin receptor site 3 per sodium channel located in the VS in *D*IV, at which α-scorpion toxins, sea anemone toxins, and related gating-modifier toxins bind ^9,10^. Consistent with this expectation from functional studies, we found a single molecule of LqhIII bound to the VS in *D*IV. The toxin binds at the extracellular end of the aqueous cleft formed by the S1-S2 and S3-S4 helical hairpins in the VS through its β2β3 loop and its C-terminal. Many conserved amino acid residues that are important for α-scorpion toxin binding and its functional effects on sodium channels are located in key positions in the toxin-receptor binding interface (Fig. 3e). These results provide convincing evidence that we have correctly identified the pharmacologically important Neurotoxin Receptor Site 3 on the native cardiac sodium channel and defined its mode of toxin binding at high resolution. α-Scorpion toxins bind to Neurotoxin Receptor Site 3 in a voltage-dependent manner, with high affinity binding to the resting state ^9,24,25^. Depolarization reduces toxin affinity and causes toxin dissociation ^9,24,25^. The bound toxin prevents the normal outward movement of the gating charges in *D*IV-S4, as measured directly from gating currents ^10^. The outward movement of the *D*IV-S4 segment correlates with fast inactivation, as measured by voltage clamp fluorometry with specifically labeled S4 residues ^12,13^. These studies highlight the importance of *D*IV-S4 and neurotoxin receptor site 3 in triggering fast inactivation. LqhIII prefers to bind to the resting state of rNa_V_1.5_C_, and strong depolarization causes dissociation in biochemical and electrophysiological studies using nM concentrations of toxin (Fig. 1) ^9,24,25^. In spite of that, the binding affinity of LqhIII for rNa_V_1.5_C_ is high enough to overcome this opposing electrostatic energy gradient when high concentrations of rNa_V_1.5_C_ and LqhIII are used to drive the binding interaction and induce gating charge transfer of *D*IV-VS into the intermediate activated state. The S4 segment of the intermediate activated *D*IV-VS is shifted two helical turns inward when compared with that of activated *D*IV-VS; however, it is not as deeply inward as in the resting state of the Na_V_Ab-VS, which suggests the toxin-induced intermediate activated state is indeed a normal intermediate state in the function of the VS. To account for the functional effects of the toxin, the toxin-modified VS must allow activation and pore opening, even while impairing fast inactivation. Trapping the *D*IV-VS in an intermediate activated state that does not trigger fast inactivation, but nevertheless is permissive for activation driven by the other three VS, would satisfy this mechanistic requirement. Thus, we propose that the toxin-induced intermediate activated state defined here is indeed the voltage-sensor trapped state that is responsible for the gating-modifier properties of the α-scorpion toxins.

Binding of LqhIII traps the voltage sensor in a unique conformation. The S4 segment is in 310 helical conformation from R1-R4, with two gating charges on each side of the HCS (Fig. 4). This conformation places the four gating charges in a straight line. R1 and R2 interact strongly with two negatively charged side chains of the ENC, whereas R3 and R4 interact strongly with two negatively charged side chains in the INC. These four ion-pair interactions bridging the HCS are unique in VS function. No other position of the primary gating charges can simultaneously form four ion-pair interactions bridging the HCS. We propose that this unique, toxin-stabilized conformation of the gating charges is the key voltage-sensor-trapped state that allows the deathstalker scorpion toxin to paralyze and kill its prey. The movement of the S4 segment implied by comparison of the structures of apo-Na_V_1.5C and LqhIII/rNa_V_1.5_C_ fits closely with the sliding-helix model of voltage sensing ^31,38,39^. Compared to the activated state, in the toxin-induced intermediate activated state, the S4 segment moves inward ~11Å and rotates slightly as 310 helix is converted to alpha helix on the intracellular side of the HCS. The gating charges exchange ion pair partners from the ENC to the INC; however, they do not move as far inward as observed in the resting state. Thus, the toxin-induced conformational change in the VS exactly follows the proposed voltage-driven movement of the S4 segment based on the sliding helix voltage-sensing model and the structures of the activated and resting states of Na_V_Ab ^31^.

Our complex structure provides an excellent model for investigating the coupling between gating charge transfer and fast inactivation. The cryo-EM structure of AaHII/Na_V_Pas-Na_V_1.7-*D*IV-VS chimera suggested that R5 and K6 of Na_V_1.7-*D*IV-VS were stabilized by interaction with the Na_V_Pas CTD. By contrast, in our fully functional LqhIII/rNa_V_1.5_C_ structure, the CTD was not observed. In fact, the CTD’s also were not observed in the high-resolution structures of human Na_V_1.2, 1.4, and 1.7 channels either ^19–21,40^. These results suggest that the CTD’s of native mammalian Na_V_ channels are disordered and/or mobile and differ substantially from the cockroach CTD in the Na_V_Pas-Na_V_1.7 chimera, whose amino acid sequence is not similar to the CTD’s of mammalian Na_V_ channels. Based on this comparison, it seems likely that the CTD plays a secondary role or a regulatory role in fast inactivation in mammalian Na_V_ channels, which may include transient interactions with the potential R5 and R6 gating charges of *D*IV-S4. Our structure gives additional insights into the structural basis for coupling of the conformational change of the intermediate activated *D*IV-VS to fast inactivation. The inward movement of the *D*IV-S4 segment and its gating charges from their activated positions propagates a voltage-driven conformational change inward to the *D*IV S4-S5 linker, forming an elbow that angles the S4-S5 linker into the cytosol. This substantial movement twists the S4-S5 linker and disturbs the conformation of the binding pocket for the IFM motif, thereby destabilizing the fast-inactivated state. In contrast, forward coupling of VS movement to fast inactivation likely involves outward movement of the S4 segment, unbending the elbow, untwisting the S4-S5 linker, and opening the receptor site for binding of the IFM motif of the fast inactivation gate. Binding of LqhIII would oppose this series of conformational events that lead to fast inactivation. Compared to the resting state, the outward movement of the *D*IV-S4 segment in the partially activated VS and the resulting loosening of the elbow bend in the *D*IV S4-S5 linker also partially open the activation gate through coupled movement of the nearby intracellular end of *D*I-S6 away from the pore axis. Our molecular dynamics analyses show that this partially open activation gate structure is still functionally closed with respect to Na^+^ conductance. Nevertheless, this conformational change may be an essential permissive movement of *D*IV that allows subsequent activation of the *D*I-*D*III VSs to drive pore opening with only partial activation of the *D*IV voltage sensor in its toxin-induced intermediate activated state. This coupling among domains may occur through the domain-swapped organization that places the *D*IV-VS adjacent to the *D*I-PM. Consistent with this idea, the *D*I-S6 segment is moved away from the axis of the pore in the toxin-induced deactivated state, similar to its movement during pore opening of Na_V_Ab. In this position, it would promote activation and pore opening of *D*II, *D*III, and *D*IV activation gate residues to give the fully open channel.

In previous work, the structure of a chimera of the cockroach sodium channel Na_V_Pas with the AaHII toxin bound was determined by cryo-EM ^26^. The functional significance of this sodium channel in the cockroach is unknown, and this chimera containing a segment of the *D*IV-VS of human Na_V_1.7 was nonfunctional; therefore, it is difficult to precisely compare this prior work to the structures we present here. Unexpectedly, AaHII bound to the Na_V_Pas chimera in two positions, one on the VS in *D*I of Na_V_Pas and one on the *D*IV-VS contributed in part by Na_V_1.7, and it was not shown whether either of these sites was functional in the chimera ^26^. In contrast, we found only a single toxin binding site, as expected from previous structure-function studies ^9,10^. Neurotoxin Receptor Site 3 identified in our study is generally similar to the AaHII binding site found in *D*IV of the AaHII/Na_V_Pas-Na_V_1.7-*D*IV-VS chimera structure ^26^, but we found an important difference in the binding poses of the two toxins. Compared with AaHII bound to the Na_V_Pas-Na_V_1.7-*D*IV-VS chimera, LqhIII bound to rNa_V_1.5_C_ is rotated ~26° downward, further away from the glycan and *D*I of the channel (Supplementary Fig. 7). This difference may reflect alteration in the position of the receptor site within the Na_V_Pas-Na_V_1.7-*D*IV-VS chimera tertiary structure caused by artifactual constraints from formation of the chimera ^26^. Alternatively, the structure of the functionally active LqhIII/rNa_V_1.5_C_ complex described here may be characteristic of the cardiac sodium channel, which has numerous distinct features compared to neuronal sodium channels like Na_V_1.7. The exact position and composition of the glycan moiety adjacent to Neurotoxin Receptor Site 3 in rNa_V_1.5_C_ may be one potentially important point of difference. Despite this difference in position of the bound toxin, the overall structures of the two toxin-bound *D*IV-VSs are remarkably similar with a RMSD of 1.41 Å over 112 residues. This striking structural similarity indicates that the *D*IV-VS’s in the two channel constructs are locked in a similar state by toxin binding at 0 mV.

Sea anemone toxins are not similar to scorpion toxins in amino acid sequence, yet they bind to Neurotoxin Receptor Site 3 and inhibit fast inactivation like α-scorpion toxins ^9,41^. Some sea anemone toxins, like Anthopleurin A and B, are highly active on cardiac sodium channels ^42,43^. It is likely that the sea anemone toxins trap the *D*IV-VS in a similar intermediate activated conformation as we have observed here for LqhIII. β-Scorpion toxins are similar in structure to α-scorpion toxins ^44^. They bind to Neurotoxin Receptor Site 4, which is located in the *D*II-VS in an analogous position to Neurotoxin Receptor Site 3 in the *D*IV-VS ^45–47^. They bind with high affinity to the activated state and trap the *D*II-VS in its outward, activated position ^45^. This mode of voltage-sensor trapping enhances activation by shifting its voltage dependence to more negative membrane potentials ^45^. When they act synergistically in scorpion venom, β-and α-scorpion toxins negatively shift the voltage dependence of activation and block fast inactivation, respectively, resulting in persistently activated sodium channels, repetitive firing, depolarization block of neuromuscular transmission, and lethal arrhythmias in the heart. A large family of cysteine-knot toxins from spiders also act as gating modifiers of voltage-gated ion channels ^48^. They bind to the VS of Na_V_, KV, and CaV channels in an analogous manner to α-scorpion toxins and either block or enhance activation of the VS. The spider toxins that enhance activation may also stabilize the unique voltage-sensor-trapped conformation of the VS that we have elucidated here, in which a linear configuration of the four primary gating charges bridges the HCS and interacts simultaneously with the ENC and INC in a high-affinity voltage-sensor-trapped complex. Thus, the toxin-bound state we have characterized here may have broad significance for voltage sensor trapping by a wide range of gating-modifier toxins from hundreds of species of spiders, scorpions, mollusks, and coelenterates, which all use this universal mechanism to immobilize their prey.

## Methods

### Electrophysiology

All experiments were performed at room temperature (21-24 °C) as described previously^21^. Human HEK293S GnTI^−^ cells were maintained and infected on cell culture plates in Dulbecco’s Modified Eagle Medium (DMEM) supplemented with 10% FBS and glutamine/penicillin/streptomycin at 37°C and 5% CO2 for electrophysiology. Unless otherwise mentioned, HEK293S GnTI^−^ cells were held at −120 mV and 100-ms pulses were applied in 10 mV increments from −120 mV to +60 mV. A P/-4 holding leak potential was set to −120 mV. Extracellular solution contained in mM: 140 NaCl, 2 CaCl_2_, 2 MgCl_2_, 10 HEPES, pH 7.4. Intracellular solution contained: 35 NaCl, 105 CsF, 10 EGTA, 10 HEPES, pH 7.4. Glass electrodes had a resistance 1.5-3 M…. Currents resulting from applied pulses were filtered at 5 kHz with a low-pass Bessel filter, and then digitized at 20 kHz. Data were acquired using an Axopatch 200B amplifier (Molecular Devices). Voltage commands were generated using Pulse 8.5 software (HEKA, Germany) and ITC18 analog-to-digital interface (Instrutech, Port Washington, NY).

### Protein expression and purification

Detailed expression and purification of rat rNa_V_1.5_C_ was described in our previous study^21^. Briefly, rNa_V_1.5_C_ was expressed in HEK293S GnTI− cells (ATCC). The protein was extracted by 1% (w/v) n-dodecyl-β-D-maltopyranoside (DDM, Anatrace) and 0.2% (w/v) cholesteryl hemisuccinate Tris salt (CHS, Anatrace) in Buffer A containing 25 mM HEPES pH = 7.4, 150 mM NaCl and 10% glycerol. After centrifugation, supernatant was agitated with anti-Flag M2-agarose resin (Sigma). Flag resin was washed in Buffer A supplemented with 0.06% glycol-diosgenin (GDN, Anatrace). Purified protein was then loaded onto a Superose-6 column (GE Healthcare) in 20 mM HEPES pH = 7.4, 150 mM NaCl and 0.06% GDN, peak fractions were concentrated to ~1 mg/ml and mixed with 50 μM LqhIII (Latoxan Laboratory) and purified FGF12b and calmodulin overnight. The mixture was then re-loaded to Superose-6 column pre-equilibrated with buffer containing 25 mM imidazole pH=6.0, 150 mM NaCl and 0.006% GDN. Finally, peak fractions were concentrated to 40 μl at 5 mg/ml.

### CryoEM grid preparation and data collection

Three microliters of purified sample were applied to glow-discharged holey gold grids (UltraAuFoil, 300 mesh, R1.2/1.3), and blotted for 2.0 − 3.5 s at 100% humidity and 4 °C before being plunged frozen in liquid ethane cooled by liquid nitrogen using a FEI Mark IV Vitrobot. All data were acquired using a Titan Krios transmission electron microscope operated at 300 kV, a Gatan K2 Summit direct detector and Gatan Quantum GIF energy filter with a slit width of 20 eV. A total of 4,222 movie stacks were automatically collected using Leginon^49^ at a nominal magnification of 130,000x with a pixel size of 0.528 Å (super-resolution mode). Defocus range was set between −1.2 and −2.8 μm. The dose rate was adjusted to 8 counts/pixel/s, and each stack was exposed for 8.4 s with 42 frames with a total dose of 60 e^−^/ Å^2^.

### Cryo-EM data processing

The movie stacks were motion-corrected with MotionCorr2^50^, binned 2-fold, and dose-weighted, yielding a pixel size of 1.056 Å. Defocus values of each aligned sum were estimated with Gctf^51^. A total of 3,805 micrographs with CTF fitted better than 6 Å were used for particle picking, and a total of 1,817,940 particles were automatically picked in RELION3.0 ^52^. After several rounds of 2D classification, 882,608 good particles were selected and subjected to one class global angular search 3D classification with an angular search step at 7.5°, at which low-pass filtered cryo-EM map of rNa_V_1.5_C_ was used as an initial model. Each of the last five iterations was further subjected to four classes of local angular search and 3D classification with an angular search step at 3.75°. After combining particles from the best 3D classes and removing duplicate particles, 570,843 particles were subjected to per-particle CTF estimation by GCTF followed by Bayesian polishing. The polished particles were subjected to last round three-class multi-reference 3D classification. The best class containing 267,595 particles was auto-refined and sharpened in Relion3.0. Local resolution was estimated by ResMap in Relion3.0. A diagram illustrating our data processing is presented in Supplementary Fig. 2.

### Model building and refinement

The structures of rat rNa_V_1.5_C_ (PDB code: 6UZ0) and LqhIII (PDB code: 1FH3) were fitted into the cryo-EM density map in Chimera^53^. The model was manually rebuilt in COOT^54^ and subsequently refined in Phenix^55^. The model vs map FSC curve was calculated by Phenix.mtrage. Statistics for cryo-EM data collection and model refinement are summarized in Supplementary Table 1.

### Molecular dynamics model

The cryo-EM structure of rNa_V_1.5_C_/LqhIII lacking *D*I-*D*II and *D*II-*D*III linkers is composed of three chains which correspond to *D*I, *D*II, and *D*III-*D*IV. The MODELLER software (ver. 9.22) was used to insert missing residues and sidechains within the polypeptide chains, followed by quick refinement using MD with simulated annealing^56^. Neutral N-and C-termini were used for the three polypeptide chains in our refined model of rNa_V_1.5_C_. N-termini from chains *D*II and *D*III-IV were acetylated, and a neutral amino terminus (−NH_2_) was used for *D*I. Neutral carboxyl groups (-COOH) were used for all C-termini. Disulfide bonds linking residues 327-342, 909-918, and 1730-1744 were included in our model of the channel as they were present in the cryo-EM structure; however, no glycans were added to the protein. Charged N-and C-termini were used for LqhIII and disulfide bonds linking residues 12-65, 16-37, 23-47, and 27-49 were included.

### Molecular dynamics simulations

Molecular models of rNa_V_1.5_C_/LqhIII with and without the toxin were prepared using the input generator, Membrane Builder^57–61^, from CHARMM-GUI^59^. The rNa_V_1.5_C_/LqhIII model was embedded in a hydrated DMPC bilayer, with approximately 150 mM NaCl. The protein was translated and rotated for membrane embedding using the PPM server^62^. The lipid bilayer was assembled using the replacement method, and solvent ions were added at random positions using a distance-based algorithm. A periodic rectangular cell with approximate dimensions of 14×14×13 nm was used, which comprised ~240,000 atoms.

The CHARMM36 all-atom force field^63–65^ was used in conjunction with the TIP3P water model ^66^. Non-bonded fixes for backbone carbonyl oxygen atoms with Na^+^ ^67^, and lipid head groups with Na^+^ ^68^ were imposed. Electrostatic interactions were calculated using the particle-mesh Ewald algorithm ^69,70^ and chemical bonds were constrained using the LINCS algorithm^71^.

The energy of the system was minimized with protein position restraints on the backbone (4000 kJ/mol/nm^2^) and side chains (2000 kJ/mol/nm^2^), as well as lipid position and dihedral restraints (1000 kJ/mol/nm^2^) using 5000 steps of steepest descent. The simulation systems were pre-equilibrated using multi-step isothermal-isovolumetric (NVT) and isothermal-isobaric (NPT) conditions for a total of 10.35 ns (see Table MDS1 for parameters). Unrestrained “production” simulations of approximately 300 ns were then generated with a 2 fs time integration step. The first 100 ns of all production simulations were considered part of equilibration based on RMSD analyses of Cα atoms (Supplementary Fig. 6) and were excluded from subsequent data analysis. Thirty independent replicas (ten of them 400 ns-long and twenty of them 300 ns-long) were generated for each system using random starting velocities, yielding a total simulation time of approximately 10.3 μs per system, of which 7.3 μs were used for analysis. The simulations were carried out using GROMACS version 2019.3 (http://www.gromacs.org).

### Molecular dynamics simulation analysis

In each snapshot of our simulations, atomic positions were translated and rotated by aligning the Cα atoms from pore transmembrane helices (S5 and S6 from all domains) of rNa_V_1.5_C_ to the initial structure produced by CHARMM-GUI. The positions of all atoms were then centered in the xy-plane by the center of mass (CoM) of pore transmembrane helices and the z-axis by the CoM of Cα atoms from the DEKA motif in the SF. After performing the spatial transformations, the z-axis of the simulation box was used as the pore axis of Na_V_1.5 and the transformed positions were used for subsequent analyses.

The axial distribution of water was computed by counting the number of water O-atoms within a cylindrical of radius 8.5 Å centered on the pore axis. The probability distribution of water was calculated for each replica by counting the number of water molecules in uniform cylindrical slices along the pore-axis and normalizing the counts by the slice with the highest number of water molecules (solvent slice). The average and SEM of the probability distribution was computed across replicas.

Pore hydration analysis indicated a dehydrated region located at the ICAG (−2.8 nm < z < −1.5 nm). The number of water molecules in the gate was counted for each frame and normalized by the total number of frames to obtain the probability distribution. The average and s.e.m. were computed across replicas.

To measure the size of the intracellular activation gate, residues at the ends of the S6 helices were selected by using a similar residue selection as Lenaeus et al. in their study of open-and closed-state Na_V_Ab structures^32^. Because rNa_V_1.5_C_ has 94% sequence similarity with hNa_V_1.5 (gap open penalty of 12, gap extension penalty of 4), a previously published multiple sequence alignment of Na_V_Ab to hNa_V_1.5 S6 helices by Boiteux et al.^72^ was used to determine the equivalent residues in rNa_V_1.5. As a result, the following residue number selections were used in rNa_V_1.5: DI: 410-413, DII: 939-942, DIII: 1469-1472, and DIV: 1771-1774. The CoM of Cα-atoms from each selection was projected onto the xy-plane and the distances between opposing S6 tails were measured (*d*_1_: DI-DIII and *d*_2_: DII-DIV).

Analyses were performed using MDTraj^73^ and molecular visualizations were rendered using Visual Molecular Dynamics^74^.

## Abbreviations

MD: Molecular dynamics
CC: central cavity
SF: selectivity filter
NaV: voltage-gated sodium channels
CoM: center of mass
SEM: standard error of the mean
RMSD: root-mean-square deviation

## Data availability

Structural Data are available from the Protein Data Bank under PDB entry ID 7K18 and EMDB entry ID EMD-2262 (https://doi.org/10.2210/pdb7K18/pdb). Reagents and additional experimental details may be obtained from.

## Acknowledgements

We thank J. D. Quispe and Quinton Beedle of the University of Washington Cryo-EM Facility for advice and technical assistance during data collection and Dr. Jin Li, Department of Pharmacology, University of Washington, for technical and editorial support. This research was supported by National Institutes of Health Research Grant R01 HL112808 (W.A.C. and N.Z.) and by the Howard Hughes Medical Institute (N.Z.), and by Canadian Institutes of Health Research grant MOP130461 (R.P.). Molecular simulations were enabled by supercomputing resources and support provided by WestGrid (*www.westgrid.ca*), SciNet (www.scinet.ca), and Compute Canada (www.computecanada.ca).

## Author Contributions

All authors designed the experiments, D. J. and L. T. carried out the cryo-EM experiments; T. M. G. carried out the electrophysiology experiments; R. B. and R. P. carried out the molecular dynamics simulations and analyses; and all authors contributed to writing and revising the manuscript.

## SUPPLEMENTARY INFORMATION

### Figures and Legends

**Supplementary Figure 1.**
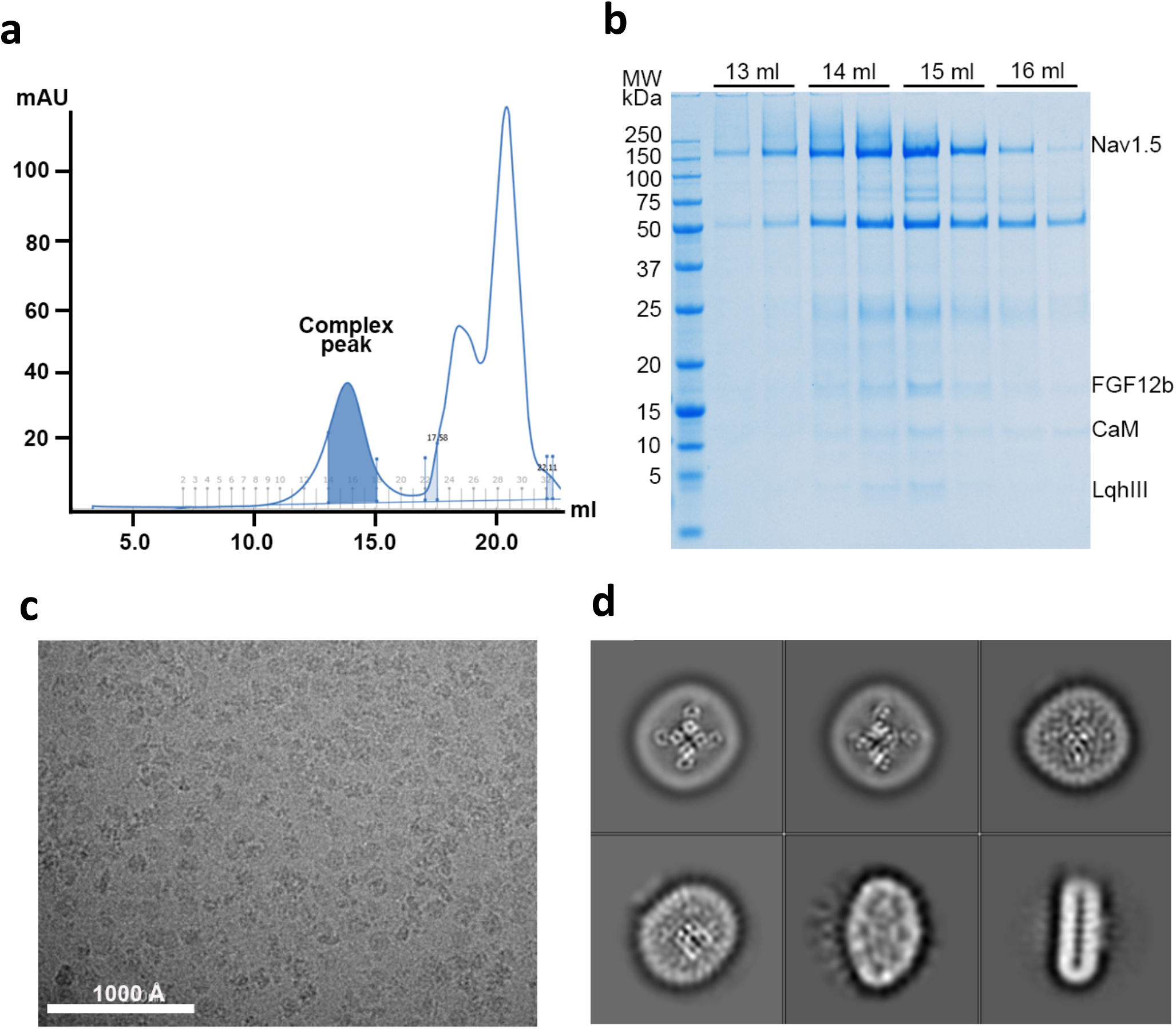
Purification and cryo-EM images of rNa_V_1.5_C_/LqhIII complex. **a**. Representative size-exclusion chromatography profile of purified rNa_V_1.5_C_/LqhIII. Peak fractions collected for cryo-EM grid preparation are shown in blue. **b.** SDS-PAGE of the SEC peak fractions stained by Coomassie blue. These results are typical of three independent preparations. **c.** Representative cryo-EM micrograph of rNa_V_1.5_C_/LqhIII sample. **d.** Selected reference-free 2D classification average.

**Supplementary Figure 2.**
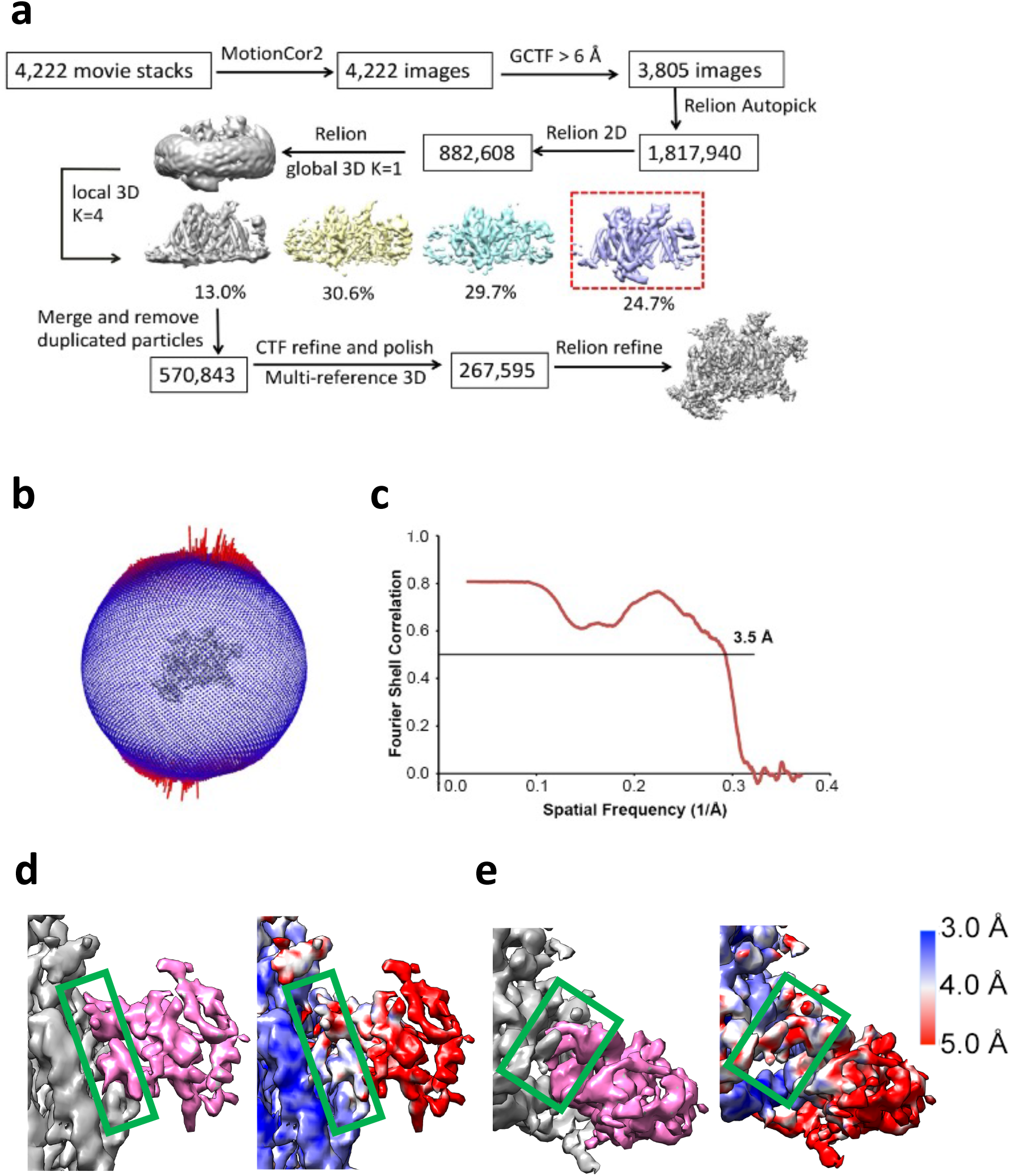
Cryo-EM data processing and resolution of the rNa_V_1.5_C_/LqhIII complex. **a**. The flowchart of the rNa_V_1.5_C_/LqhIII data processing. **b**. Particle angular distribution of the final reconstruction. **c**. The Fourier Shell Correlation curve of the model versus the map used for model refinement. **d.** Expanded side view of the local resolution of the rNa_V_1.5_C_/LqhIII interface. e. Expanded top view of the local resolution of the rNa_V_1.5_C_/LqhIII interface.

**Supplementary Figure 3.**
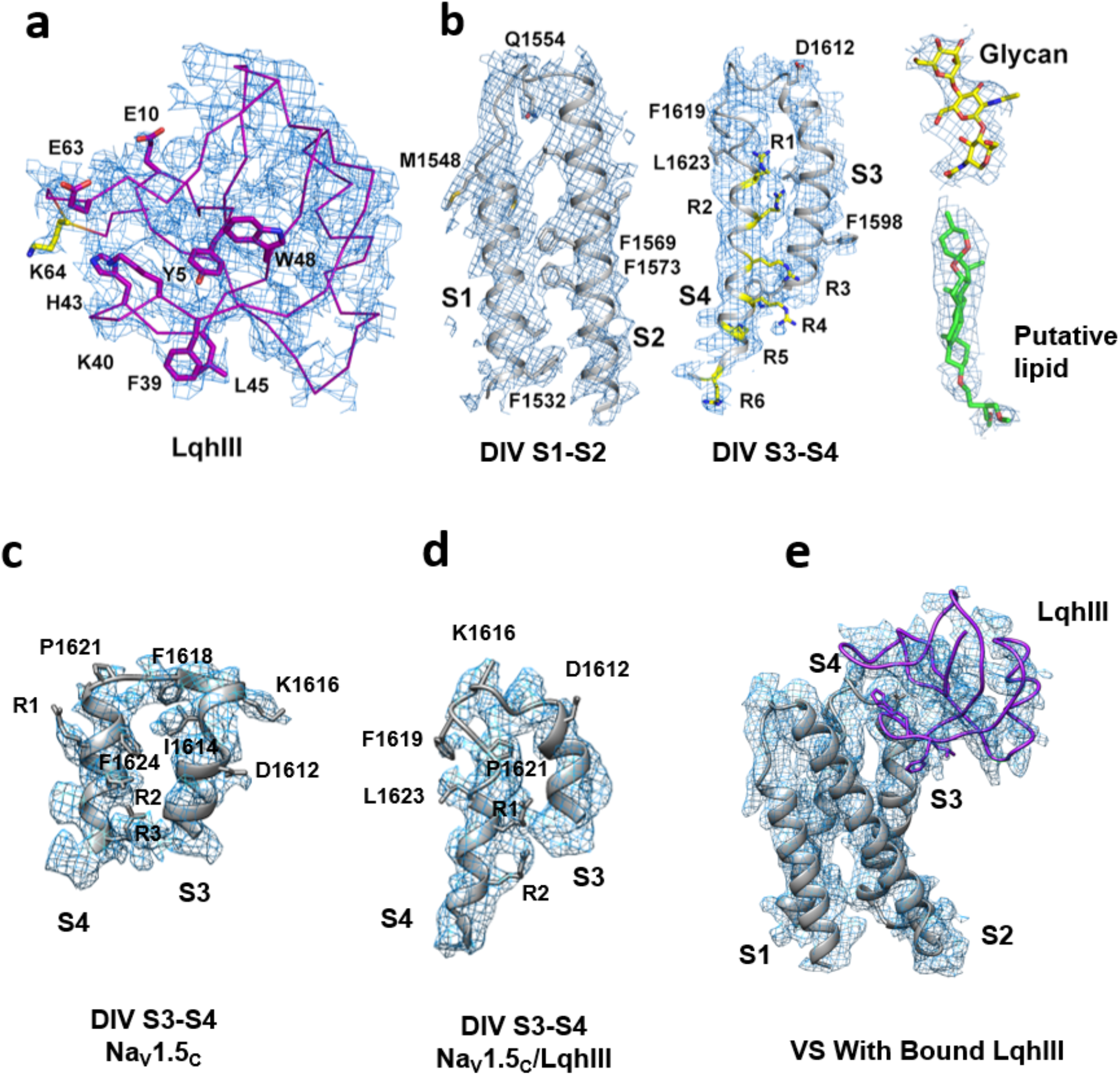
Representative EM map for key extracellular components of the rNa_V_1.5_C_/LqhIII complex. **a**. The EM density for LqhIII. **b**. The EM density for *D*IV S1-S2, *D*IV S3-S4, N329 linked glycan, and a putative lipid (modeled as the detergent GDN in green sticks) inside the activation gate, respectively. Key residues are shown in sticks. **c**. EM density for the S3-S4 linker in rNa_V_1.5_C_. **d**. EM density for the S3-S4 linker in Na_V_1.5_c_/LqhIII. **e**. EM density for the complex of Na_V_1.5_C_/S3-S4 and LqhIII.

**Supplementary Figure 4.**
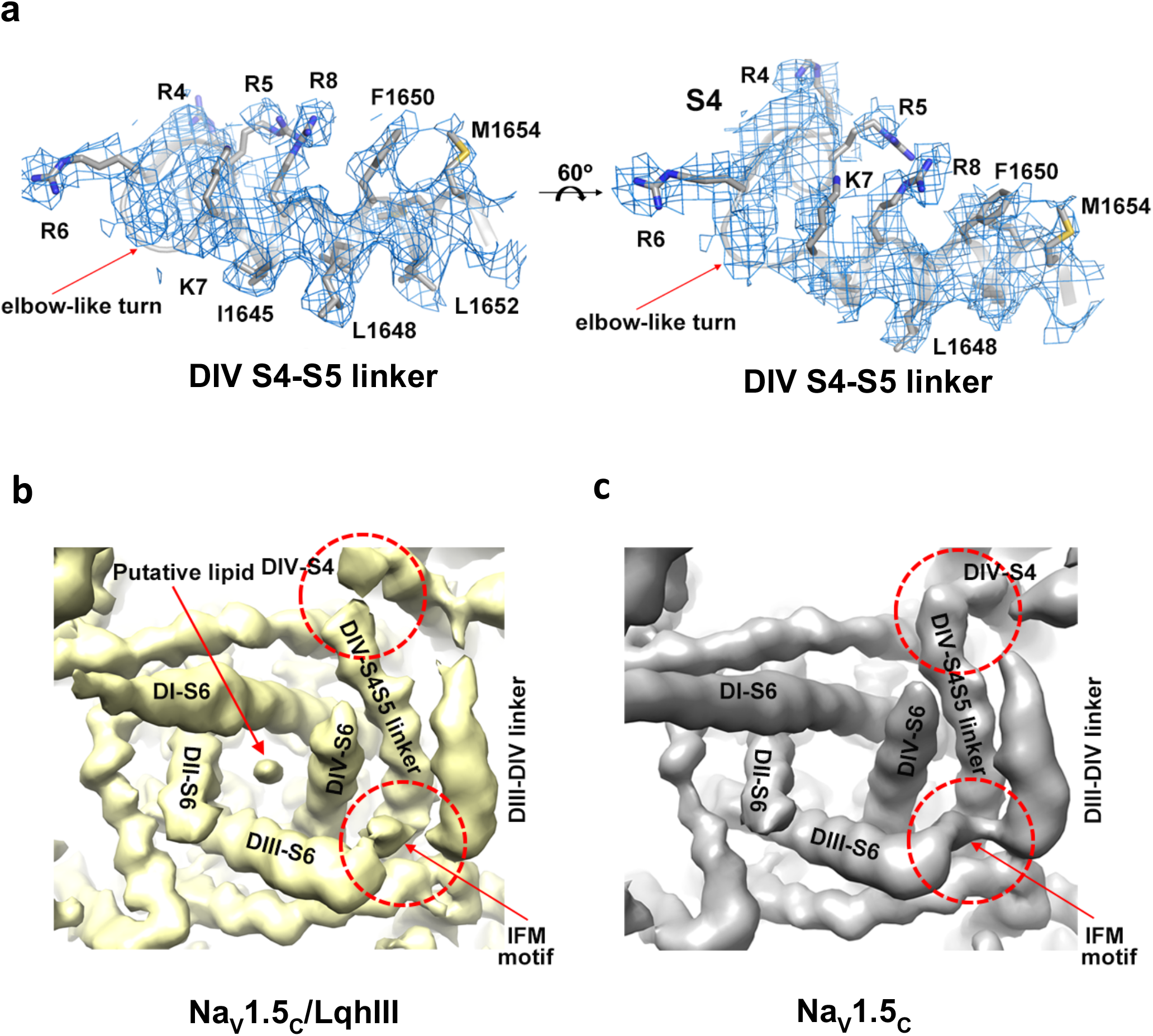
Representative EM map for key intracellular components of the rNa_V_1.5_C_/LqhIII complex. **a**. EM density for the rNa_V_1.5_C_/LqhIII *D*IV S4-S5 linker. (**b-c**) The unsharpened EM density comparison between rNa_V_1.5_C_/LqhIII (yellow) and rNa_V_1.5_C_ without toxin (grey) in a bottom (intracellular) view of the intracellular activation gate and inactivation gate. Red dash circles indicated the areas of conformational change in the two maps.

**Supplementary Figure 5.**
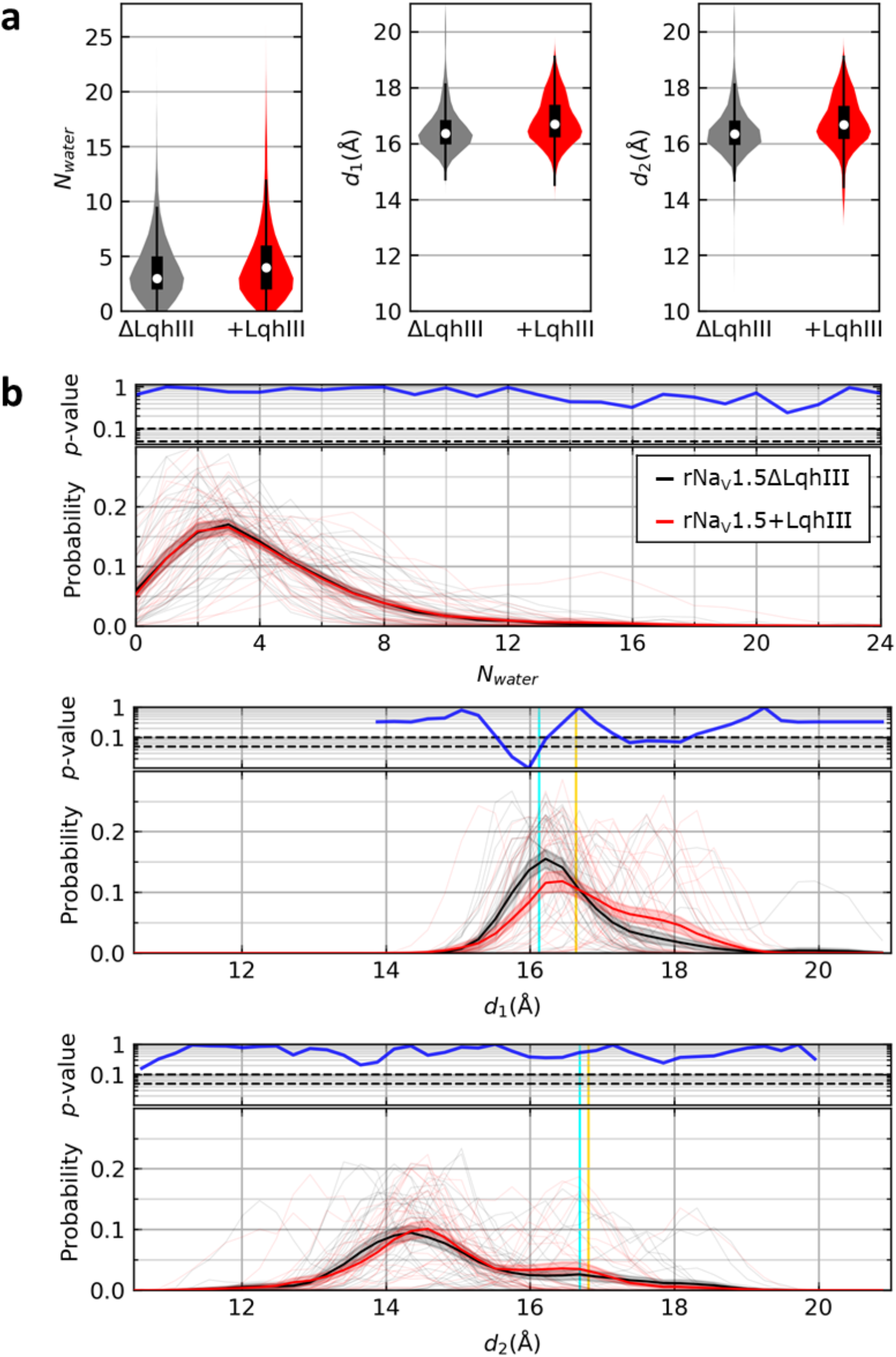
Significance of changes in size and hydration of the intracellular activation gate. **a.** Differences in mean values of hydration and diagonal distances of the intracellular activation gate due to toxin binding. Violin plots of means and s.e.m. Left, the number of water molecules in the intracellular gate (*N*water). Middle, *d_1_* (*D*I-*D*III). Right, *d_2_* (*D*II-*D*IV) for simulations of rNa_V_1.5_C_ with (red) or without (grey) LqhIII. The median (white dot), quartiles (thick black line), and interquartile ranges (thin black line) are shown. *p*-values were calculated from a two-sided Welch’s t-test which indicated significant differences in the means of the distributions between simulations: Left: *N*water = 3.958 ± 0.017 for Na_V_1.5C; *N*water = 4.307 ± 0.019 for rNa_V_1.5_C_/LqhIII, p=2×10^−42^. Middle: *d*_1_ = 16.520 ± 0.005 for rNa_V_1.5_C_; *d*_1_ = 16.810 ± 0.001 for rNa_V_1.5_C_/LqhIII; p=0.001. Right: *d*_2_ = 16.465 ± 0.005 for rNa_V_1.5_C_; *d*_2_ = 16.744 ± 0.005 for rNa_V_1.5_C_/LqhIII; p=7×10^−300^. Mean and s.e.m. Time frames spread across 30 independent simulations were used for significance testing (n=29,026 frames for rNa_V_1.5_C_ and n=29,899 frames for rNa_V_1.5_C_/LqhIII). **b.** Changes in hydration and diagonal distances of the intracellular activation gate due to LqhIII binding. Top. *p*-values calculated from a two-sided Welch’s t-test for the probability of *N*water in n=30 independent MD simulations of the cryo-EM rNa_V_1.5_C_ structure with or without LqhIII. Toxin-binding insignificantly changes the hydration of the gate. Significance levels of 0.1 and 0.05 are indicated as black and dashed lines. Bottom. The thick line with shading is the average probability distribution of *N_water_* and the thin and transparent lines are distributions from 30 independent replicas for simulations with (red) or without (black) LqhIII. Shading represents s.e.m. In the presence of the toxin, *d_1_* is significantly larger, as indicated by both a significant decrease in the probability of *d_1_* = 15.8 Å to 16.2 Å ([*d_1_* (Å)*, p*] = {(15.8, 2×10^−2^), (16.0, 1×10^−2^), (16.2, 9×10^−2^)}) and increase in the probability of *d_1_* = 17.4 Å to 18.0 Å ([*d_1_* (Å)*, p*] = {(17.4, 7×10^−2^), (17.6, 8×10^−2^), (17.9, 8×10^−2^), (18.0, 8×10^−2^)}). The *p*-values and average probability distribution of diagonal distance, *d_2_*, between S6 helix tails of *D*II and *D*IV. No significant change is observed. *d_1_* and *d_2_* of the reference cryo-EM structures of rNa_V_1.5_C_ (PDB ID: 6UZ3; cyan) and rNa_V_1.5_C_/LqhIII (yellow) are indicated by colored vertical lines.

**Supplementary Figure 6.**
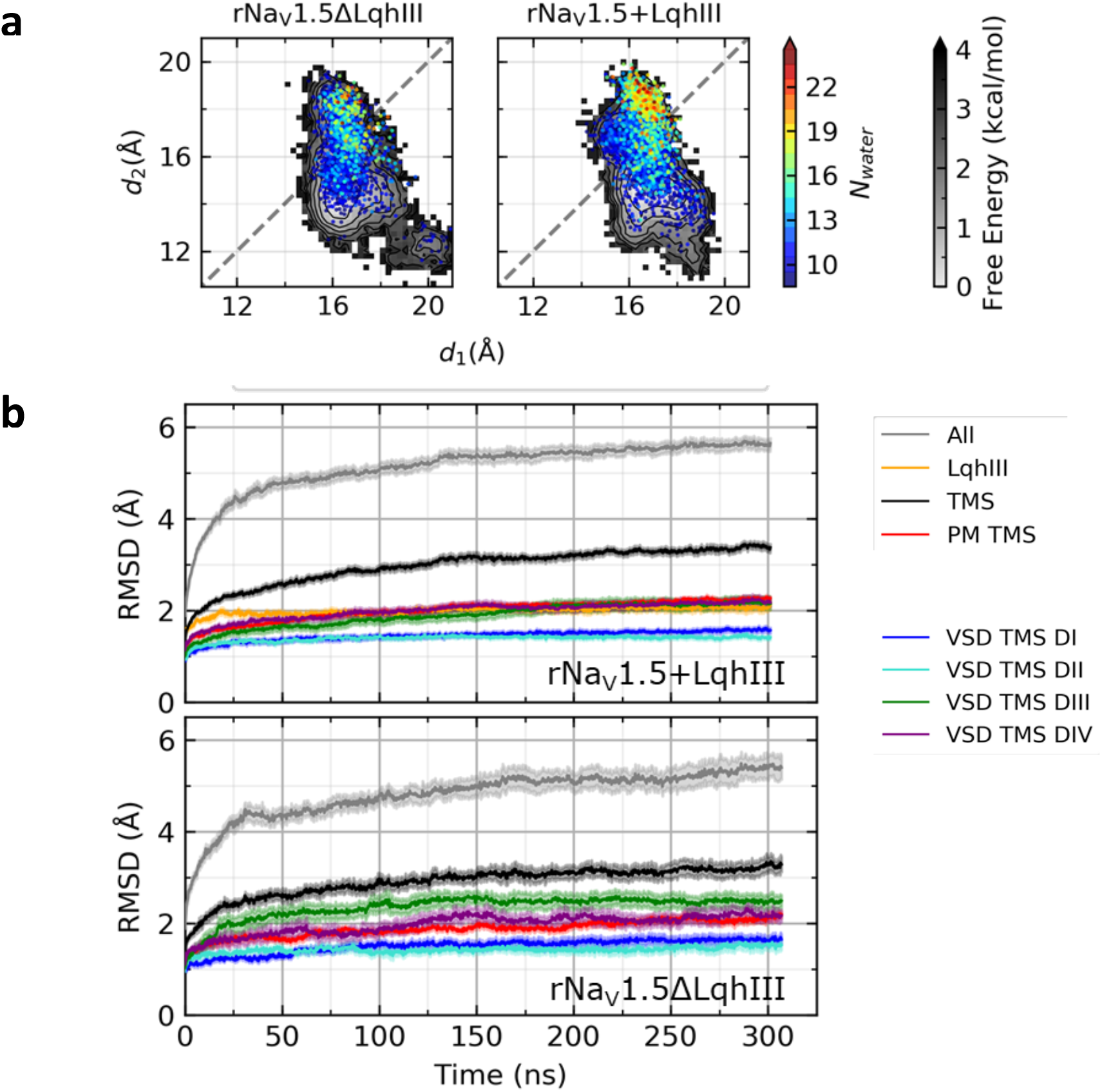
Hydration and time evolution of diameter in the intracellular activation gate. **a.**Hydration of the intracellular activation gate at dilated conformations. Free energy distribution of diagonal distances (grey-scale) between S6 helix tails (*d_1_*: DI-DIII, *d_2_*: DII-DIV); same as Figure 6. *d_1_* and *d_2_* conformations corresponding to *N_water_* ≥ 9 are shown as circles colored according to *N_water_*. Conformations with *N_water_* ≥ 15 are most often found when *d_2_* ≥ *d_1_* > 16 Å. **b.**Time evolution of the protein root-mean-square deviation (RMSD). The average RMSD was calculated over time relative to the initial cryo-EM rNa_V_1.5_C_/LqhIII structure for simulations with (top) or without (bottom) the toxin. The RMSD of backbone Cα atoms for all residues (grey), LqhIII only (orange), transmembrane segments (TMS) of rNa_V_1.5_C_ (black), as well as TM segments from the pore module (PM; red) and voltage sensing domains (VS) from *D*I (blue), *D*II (cyan), *D*III (green), and *D*IV (purple) are shown. Based on PM, TMS, and LqhIII RMSD values, simulations equilibrate at approximately 100 ns. Time frames spread across 30 independent simulations of rNa_V_1.5_C_/LqhIII with or without the toxin were used for analyses (n=29,026 frames for rNa_V_1.5_C_ and n=29,899 frames for rNa_V_1.5_C_/LqhIII).

**Supplementary Figure 7.**
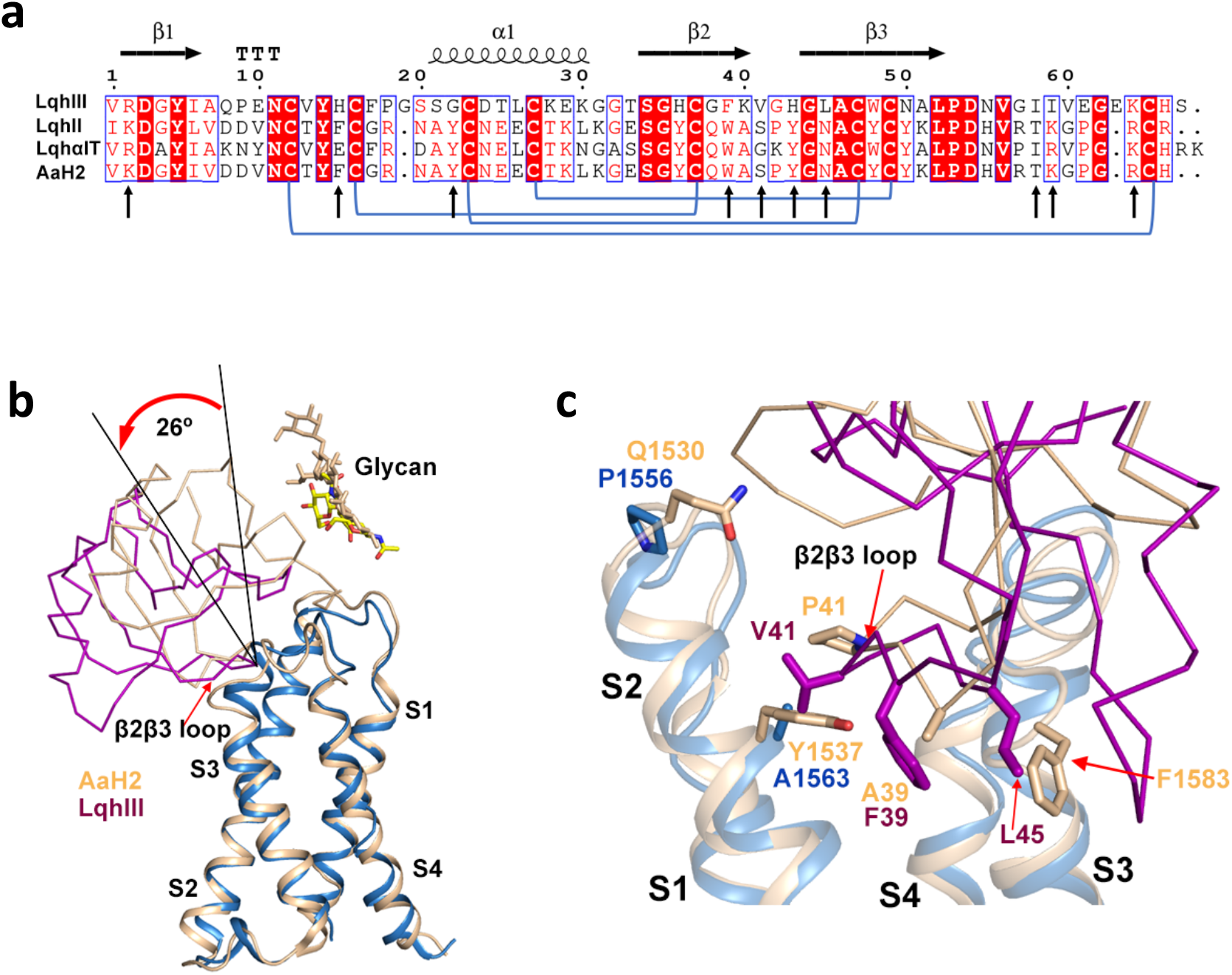
Comparison of the structure of the Na_V_1.7/Na_V_Pas/AaHII complex to the structure of the rNa_V_1.5_C_/LqhIII complex. **a**. Sequence alignment among α-scorpion toxins. Disulfide-bond forming Cys were connected by blue lines. Key residues from site-directed mutagenesis studies were indicated with black arrows. **b**. Superposition of rNa_V_1.5_C_ *D*IV-VS/LqhIII and Na_V_1.7 *D*IV-VS/AaHII. Na_V_1.7 *D*IV-VS/AaHII was colored in wheat. **c**. Comparison of the binding positions in the rNa_V_1.5_C_/LqhIII complex (purple) and the Na_V_Pas-Na_V_1.7-*D*IV/AaHII complex (wheat). Key residues shown in sticks.

**Table S1.** Cryo-EM data collection, refinement and validation statistics.

|  |  |
| --- | --- |
|  | rNa<sub>v</sub>1.5<sub>c</sub>/LqhIII<br>(EMDB code: EMD-22621)<br>(PDB code: 7K18) |
| **Data collection and processing** |  |
| Magnification | 130,000 |
| Voltage (kV) | 300 |
| Electron exposure (e-/Å<sup>2</sup>) | 60 |
| Defocus range (μm) | 1.2 – 2.8 |
| Pixel size (Å) | 1.056 |
| Symmetry imposed | C1 |
| Initial particle images (no.) | 1,817,940 |
| Final particle images (no.) | 267,595 |
| Map resolution (Å) | 3.34 |
| FSC threshold | 0.143 |
| Map resolution range (Å) | 3.0 – 5.0 |
| **Refinement** |  |
| Initial model used (EMDB code) |  |
| Model resolution (Å) | 3.5 |
| FSC threshold | 0.5 |
| Model resolution range (Å) | 3.0 – 5.0 |
| Map sharpening *B* factor (Å<sup>2</sup>) | -86 |
| Model composition |  |
| Non-hydrogen atoms | 9973 |
| Protein residues | 1195 |
| Ligand | 25 |
| *B* factors (Å<sup>2</sup>) |  |
| Protein | 59.0 |
| Ligand | 53.1 |
| R.m.s. deviations |  |
| Bonds (Å) | 0.004 |
| Angles (°) | 0.758 |
| Validations |  |
| MolProbity score | 2.75 |
| Poor rotamers (%) | 0.8 |
| Ramachandran plot |  |
| Favored (%) | 93.99 |
| Allowed (%) | 5.76 |
| Disallowed (%) | 0.25 |

**Table S2.** Molecular Dynamics Simulation Parameters. MD parameters used for each pre-equilibration and production step. The ensemble (ENS), time integration step, and total time/replica are indicated. Berendsen (B) (1) or Nosé-Hoover (NH) (2, 3) thermostats and B or Parrinello-Rahman (PR) (4, 5) barostats with their respective coupling times constants (CTC) are shown. Protein backbone and sidechain position restraints as well as lipid position and dihedral restraints are also specified.

| Step | ENS | Time Step<br>(fs) | Total Time<br>(ns) | Temperature (T) |  |  | Pressure (P) |  |  | Restraints<br>(kJ/mol/nm<sup>2</sup>) |  |  |  |
| --- | --- | --- | --- | --- | --- | --- | --- | --- | --- | --- | --- | --- | --- |
|  |  |  |  | T<br>(K) | Thermo-<br>stat | T<br>CTC | P<br>(bar) | Baro-<br>stat | P<br>CTC | Protein<br>Backbone | Protein<br>Sidechain | Lipid<br>Position | Lipid<br>Dihedral |
| 1 | NVT | 1 | 0.025 | 300 | B | 1 ps | --- | --- | --- | 4000 | 2000 | 1000 | 1000 |
| 2 | NVT | 1 | 0.05 | 300 | B | 1 ps | --- | --- | --- | 2000 | 1000 | 1000 | 400 |
| 3 | NPT | 2 | 10 | 300 | B | 1 ps | 1 | B | 5 ps | 1000 | 500 | 400 | 200 |
| 4 | NPT | 2 | 0.1 | 300 | B | 1 ps | 1 | B | 5 ps | 500 | 200 | 200 | 200 |
| 5 | NPT | 2 | 0.1 | 300 | B | 1 ps | 1 | B | 5 ps | 200 | 50 | 40 | 100 |
| 6 | NPT | 2 | 0.1 | 300 | B | 1 ps | 1 | B | 5 ps | 50 | 0 | 0 | 0 |
| 7 | NPT | 2 | 300-400 | 300 | NH | 1 ps | 1 | PR | 5 ps | 0 | 0 | 0 | 0 |

